# Multiomic Spatial Imaging Assay (MSIA) – A High-plex *In Situ* Detection Method for mRNAs, Proteins and Protein-Protein Interactions using Manual And Semi-Automated Workflows

**DOI:** 10.64898/2026.02.26.708124

**Authors:** Chengxin Zhou, Alvason Li, Ji Zhang, Yifan Wang, Yuanyuan Liu, Pehr Williamson, Ge-Ah Kim, Sonali A Deshpande, Miao Yuan, Suganya Chandrababu, Lina Duan, Ching-Wei Chang, Li-Chong Wang, Maithreyan Srinivasan

## Abstract

We report Multiomics Spatial Image Analysis (MSIA), a suite of technologies that include streamlined manual and semi-automated workflows to image 100s of RNAs from fresh frozen and formalin-fixed paraffin-embedded (FFPE) samples, contextualize RNAs with select protein markers including imaging synaptic protein interactions, a data analysis pipeline that includes corrections for chromatic aberration, algorithms for image stitching, deep learning models for image analysis (cell segmentation and RNA dot detection using hundreds of RNAscope images) from multiple stain-image cycles and proof of concept for a language model trained on extensive Parkinson’s Disease (PD) literature. We used the language model to identify potential new biomarkers that were confirmed in MPTP-treated mouse model of PD. Furthermore, by incorporating more stain-image cycles alongside an additional fluorescent channel and advanced error-correcting decoding schemes, the MSIA workflow demonstrates the scalability to potentially quantify over 1,000 genes.

## Introduction

In parallel with advances in genomics, spatial multiomics technologies have emerged as powerful tools to interrogate tissues at single-cell/subcellular resolution, integrating mRNA and protein data. These approaches, whether sequencing- or imaging-based, offer unprecedented throughput and spatial context for biological samples [1]. Many of these approaches are available as commercial solutions that cost hundreds of thousands of dollars making these technologies out of reach for many researchers.

Here we report a prototype assay named Multiomics Spatial Imaging Analysis (**MSIA**), which utilizes manual and semi-automated Leica Bond Rx workflows combined with conventional fluorescent microscopy without the need for expensive instrumentation.

MSIA is built on the foundations of the well-established RNAscope, the first commercial platform that standardized spatial RNA measurements. RNAscope relies on highly specific and sensitive in situ hybridization chemistry, with probe design and signal amplification steps that enable quantitative measurements of gene expression at single-molecule resolution. In this study, we report the interrogation of 100 genes across <u>only</u> <u>two stain–image cycles</u> in both fresh-frozen and formalin-fixed, paraffin-embedded (FFPE) tissues. We first used MSIA to evaluate the transcriptomic changes in brain aging process using wild type mice of different ages and compared our data with similar data collected and published by other spatial transcriptomic technology developers. In addition, to highlight the reduced cost and analytic complexity, we deployed the technology on smaller, hypothesis-driven gene sets relevant for Parkinson’s disease biomarker discovery.

Parkinson’s disease (PD), the second most prevalent neurodegenerative disorder after Alzheimer’s disease, is marked by progressive deterioration of motor and non-motor functions. This decline arises largely from decreased dopamine levels in the substantia nigra and widespread accumulation of misfolded α-synuclein aggregates (Lewy bodies) throughout the brain [2]. Although significant research has focused on the molecular underpinnings of PD, its clinical heterogeneity and incomplete mechanistic understanding continue to pose substantial challenges for early diagnosis, patient stratification, and therapeutic development. Genetic predisposition, both in terms of specific causal variants and common variants contributing to polygenic risk, accounts for 22–40% of PD heritability [2–4]. However, reduced penetrance in most monogenic forms highlights the essential interplay between genetic and environmental factors.

Numerous studies further implicate gene expression changes—particularly alterations in mRNA levels at early disease stages—in disease onset and progression [5]. Yet, capturing these expressions shifts before the emergence of microscopic Lewy bodies and other prodromal markers that suggest pace of disease progression remain difficult. Early identification of such changes would profoundly enhance strategies for prevention, intervention, and biomarker discovery. The main objective of this study is to report on the development of a complete MSIA workflow that includes assay chemistry and software analysis packages. We show evidence that this method can be deployed on diverse sample types, report the identification of potential new mRNA biomarkers using language models and their verification on a MPTP-induced PD mouse model.

Several innovations as part of this study are described

1. Utilized a novel combinatorial optical barcoding method based on the well-established RNAscope chemistry [6, 7] to develop MSIA. We profiled 100 differentially expressed genes implicated in aging and validated the technology using fresh-frozen and FFPE brain sections collected from wild-type mice of 4 age groups: 4, 21, 52, and 90 weeks. To further validate the method, we used 53 established genes implicated in PD and performed MSIA. MSIA can decode up to 100 genes in only two staining/imaging cycles and does not require expensive instrumentation or error correction methods. It is performed manually, and the slides can be imaged using a conventional fluorescence microscope, rendering them more feasible for many translational research settings.
2. Autofluorescence removal
3. To add orthogonal cell-cell interaction confirmation and proteins, MSIA was executed in 3 cycles, with 50-100 RNA targets in cycles 1 & 2 followed by an additional cycle to detect critical protein targets, thereby validating and expanding our transcriptional data with protein-level insights that collectively mark different neuronal/non-neuronal cells and synaptic neurexin-neuroligin signaling complexes.
4. Chromatic aberration correction
5. Deep-learning (DL)-based cell segmentation algorithms were trained using manually annotated ground-truth data from internal archived images.
6. In-house dot detection algorithms using deep learning methods on 100s of archival RNAscope images from mouse brain tissues.
7. To further advance gene discovery, we harnessed a language model trained on extensive PD literature to identify underexplored, low-expression genes potentially implicated in disease onset by assigning high dimensional vector embeddings to words in literature text in a way that retains the semantic and syntactic relationships.
8. Predicted candidate genes were then tested in PD animal models establishing the connection between these newly uncovered transcripts and established PD pathophysiology.

Altogether, this integrated approach—spanning multiomic spatial analysis, computational image processing, and literature-mining large language models—promises to deepen our understanding of PD pathogenesis and accelerate the search for novel biomarkers and therapeutic targets.

## Materials and Method

### Materials

C57Bl/6J mice 8-weeks old wild type. (Jackson Laboratory, Maine, US). Food steamer (Model: 5712; Oster)

ACD EZ hybridization system, MSIA ISH probes, RNAscope tissue pretreatment kits, and RNAscope HiPlex v2 assay kits (Bio-techne)

Zeiss Axio Imager Z2 microscope with XY Piezo stage, Axiocam 712 mono camera, and Colibri7 light source (Zeiss)

Filter sets: Filter Set 110 HE LED (Zeiss), Dylight 550 Filter Set (Chroma) and Dylight 650 Filter Set (Chroma)

Code availability: ACDnet code and model used to generate the main figures of this paper can be available for academic use through GitHub repository upon request: https://github.com/alvason/ACDnet_RNAdot_100plex

### Methods

A total of 16 C57BL/6J mice (n=8 female and n=8 male), 8-10 weeks old were used in the study. In the MPTP model, the bilateral dopaminergic neuronal death of the substantia nigra and dopamine depletion of the striatum was created by repeated intraperitoneal injection of 20 mg/kg MPTP. This MPTP regimen generates early-symptomatic, motor-predominant PD state that typically results in ∼50% loss in substantia nigra [2]. Mice were injected 4 times at 12-hour intervals for two consecutive days (Day 0 and 1) to a total amount of 80mg/kg. In another group, 8 litter-mate mice (4 male and 4 female) were injected with saline and used as the vehicle control for the treated group. These mice were subjected to fine motor kinematic gait analysis on day 7. On day 8, mice were euthanized, and brain samples were collected and prepared into FFPE blocks (**Supplementary Figure. S1.A**).

### Tissue collection

Mouse brain was sectioned from fixed-frozen and FFPE mouse brains from 1 month, 5 months, 12months and 24 months old mice. All sectioned mouse brain samples were acquired from Acepix Biosciences (Union City, CA). C56BL/6J mice of different ages were acquired from Jackson Laboratory (Bar Harbor, ME) via a custom mouse aging program. Mice were subjected to transcardiac perfusion with 4% PFA before decapitation. Whole mouse brains were collected and further fixed in 10% NBF for 24hrs. After dehydration with 30% sucrose, some tissues were then embedded in Optimal Cutting Temperature (OCT) compound (Sakura, USA), snap-frozen in liquid nitrogen, and sectioned coronally from the brain block with a cryotome. Brain sections were post-fixed in 10% NBF and dehydrated in ethanol before the assay.

For FFPE processed brains, freshly dissected mouse brains were then trimmed, placed into a tissue cassette, and embedded in paraffin using an automated processor.

### Target probe selections, synthesis, and formulation

A collection of hundreds of mouse neuro-specific and PD-related mouse genes were selected as the targets of MSIA transcriptomic profiling. These genes were examined for their expression levels, spatial expression pattern in the mouse brains, and specificity to intended brain cell types using single-cell RNAseq datasets from public scRNA-Seq data deposit. RNAscope Multiplex and HiPlex assays were performed on the coronal mouse brain sections to further verify the spatial expression patterns of some high-expression genes, in efforts to avoid any spatial crowding in the final FISH images, which could otherwise complicate the decoding process.

All target probes were designed by ACD probe design team and were aligned to different transcript databases using BLAST program [8] to ensure no potential cross-reactivity to other RNA species. Probe design pipeline was modified to accommodate 100s of genes that satisfy codebook rules (**Table-2** and **Figure.1**) All DNA oligo probes are formulated at 10uM concentration and were pooled in ACD probe diluent at 1:500 dilution. 100 target probes were designed and synthesized for the first cycle and another set of 100 target probes for the second cycle of the assay.

### Barcode scheme

The assay is built on a codebook that is composed of 10 fundamental barcodes. Each fundamental barcode has 5 bits of information including two fluorescence signals (e.g. ‘10100’ has a AF488 signal and a Dy594 signal). To ensure high fidelity in the ability to detect expressed genes, sparse barcoding methods that are expected to result in less dot crowding were used. A pool of 5 unique fluorophores is used to generate various combinations of two fluorophores, which include: AF488, Dy550, Dy594, Dy650, and AF750. The resultant 5-bit barcodes are listed below (**Table.2**):

To reach the 100plex analytic capacity, we expanded the 5-bit primary barcodes to create a new codebook consisting of 100 ‘10-bit’ target barcodes. This is accomplished by running two barcoding cycles, where the primary 5-bit barcode is randomly paired with another 5-bit barcode from the list above. Each 10-bit barcode is unique in the codebook and can be assigned to a gene or protein target. A list of 100 CNS-specific targets, each paired with a uniquely designated 10-bit barcode, is available in the Supplementary Table.1.

All barcodes have 10 bits with a Hamming weight of 4 (2 fluorophore bits out of 5 bits + 2 stain and image cycles). Due to the nature of the assay, the barcoding method has no temporal dependency. Because each barcoding cycle is chemically independent from others, and thus can be performed in any temporal sequence, target barcodes need not follow in a sequential order. Moreover, in contrast to MERScope, the barcoding system has curated a collection of high-fidelity barcodes that are best suited for conventional epifluorescence microscopes and our proprietary deep-learning-based decoding method. This strategy led to an accurate decoding outcome with a low error rate (non-assigned barcode ratio: 0.3∼0.6%). This is extremely beneficial, as it removes the potential need for an error-correction mechanism, which requires a larger minimum Hamming Distance between utilized barcodes, hence more cycles and/or fluorescence channels, to achieve the same number of usable barcodes. We therefore are able to use a minimum Hamming Distance of only 2 (which does not allow error correction but allows for more efficient use of barcodes) and a fixed Hamming Weight of 4 for our barcodes, as listed in **Table 2**.

**Table 1:**
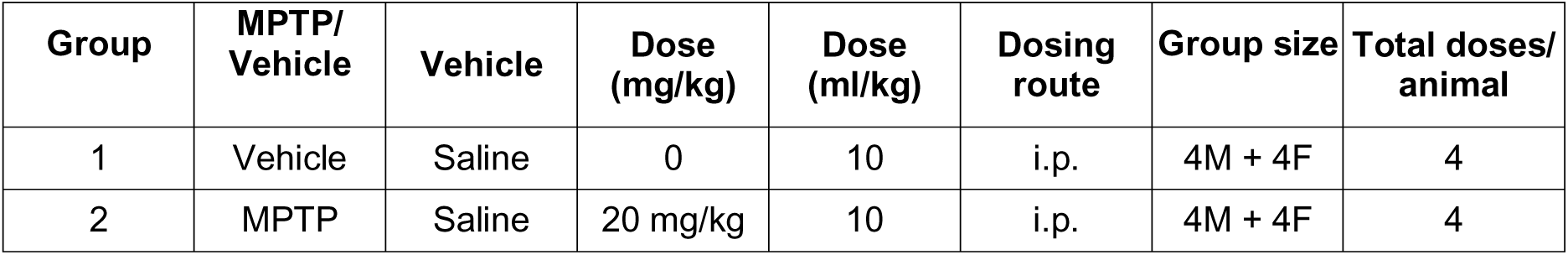
Animal assignment and drug administration scheme. 1- methyl-4-phenyl-1,2,3,6-tetrahydropyridine (**MPTP) Mouse Model**

**Table 2:** The list of 10 primary, combinatorial barcodes used in 100plex MSIA. Each barcode has 5 bits and contains two fluorescence signals.

| S.N | fluor bits | 488 | 550 | 594 | 650 | 750 |
| --- | --- | --- | --- | --- | --- | --- |
| *a* | 2-fluorophore | 0 | 0 | 0 | 1 | 1 |
| *b* | 2-fluorophore | 0 | 0 | 1 | 0 | 1 |
| *c* | 2-fluorophore | 0 | 0 | 1 | 1 | 0 |
| *d* | 2-fluorophore | 0 | 1 | 0 | 0 | 1 |
| *e* | 2-fluorophore | 0 | 1 | 0 | 1 | 0 |
| *f* | 2-fluorophore | 0 | 1 | 1 | 0 | 0 |
| *g* | 2-fluorophore | 1 | 0 | 0 | 0 | 1 |
| *h* | 2-fluorophore | 1 | 0 | 0 | 1 | 0 |
| *i* | 2-fluorophore | 1 | 0 | 1 | 0 | 0 |
| *j* | 2-fluorophore | 1 | 1 | 0 | 0 | 0 |

**Table 3:**
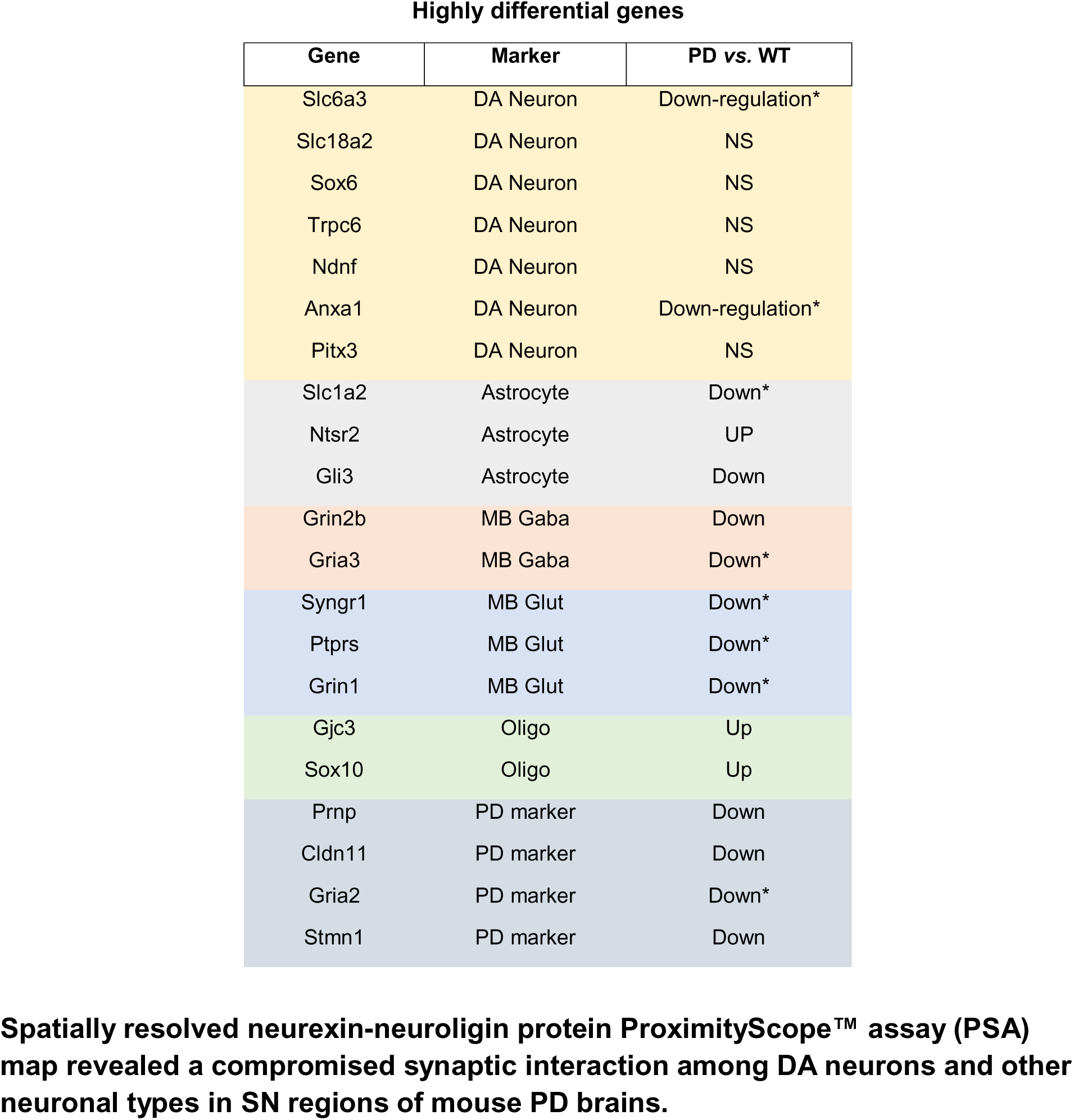
MSIA assay unveiled mRNA expressional changes in MPTP mouse brains. (* results consistent with prior studies.; NS: not significantly changed.)

### RNA molecule labeling

To encode the RNA molecules with corresponding barcodes, each RNA transcript is first hybridized to a specific double-Z target probe. The target probe is then annealed to 3 tiers of branching DNA oligos (i.e. AMP1, AMP2, and AMP3) to form the signal generation complex (SGC). Each of the 5-bit barcodes uses a unique SGC that has been characterized for no detectable cross talk [6, 9]. The third AMP oligo contains a unique binding sequence that is complementary to the corresponding fluorophore-conjugated oligos (**Label Probe, or LP**). The development of signal generating configuration is completed upon incubating the branched DNA structure with its corresponding label probes. A large body of previous work has established no detectable crosstalk between the different branched DNA structures [7]. For each RNA species, the branch DNA structure expected to hybridize to is now annealed with its complementary label probes that are conjugated with either fluorophore A or fluorophore B (50%:50% LP-A /LP-B in the final solution), as pre-determined in the codebook (**Table.2**). This allows the RNA molecule to emit two fluorescence signals and therefore display a unique 5-bit barcode in the acquired multi-spectral image. All 100 RNA targets included in the panel were encoded twice through two independent barcoding cycles. The first cycle of barcoding provides the first 5 bits of the barcode, and the second cycle provides the last 5 bits of the barcode. In each cycle, all 100 genes were <u>simultaneously</u> labeled with their corresponding 5-bit barcodes, which does not follow any temporal order of barcoding (**Figure 1**). This is an important distinction with other methods such as Xenium and MERScope [10, 11] In addition, the two stain-image cycles are independent of each other. Each barcoding cycle can be performed either first or last. After the first cycle, any prior branching DNA structure is stripped off from the RNA molecules with HiPlexUp reagent [12], and the RNA molecules were subject to another round of signal generation beginning with a new target probe hybridization. It should be noted that this approach can be used to perform additional rounds of probe hybridization and signal generation to increase throughput. This assay has been optimized both as a streamlined manual as well as semi-automated workflows using the Leica BondRx autostainer. Both approaches resulted in excellent positive signals and a very clean background.

**Figure 1.**
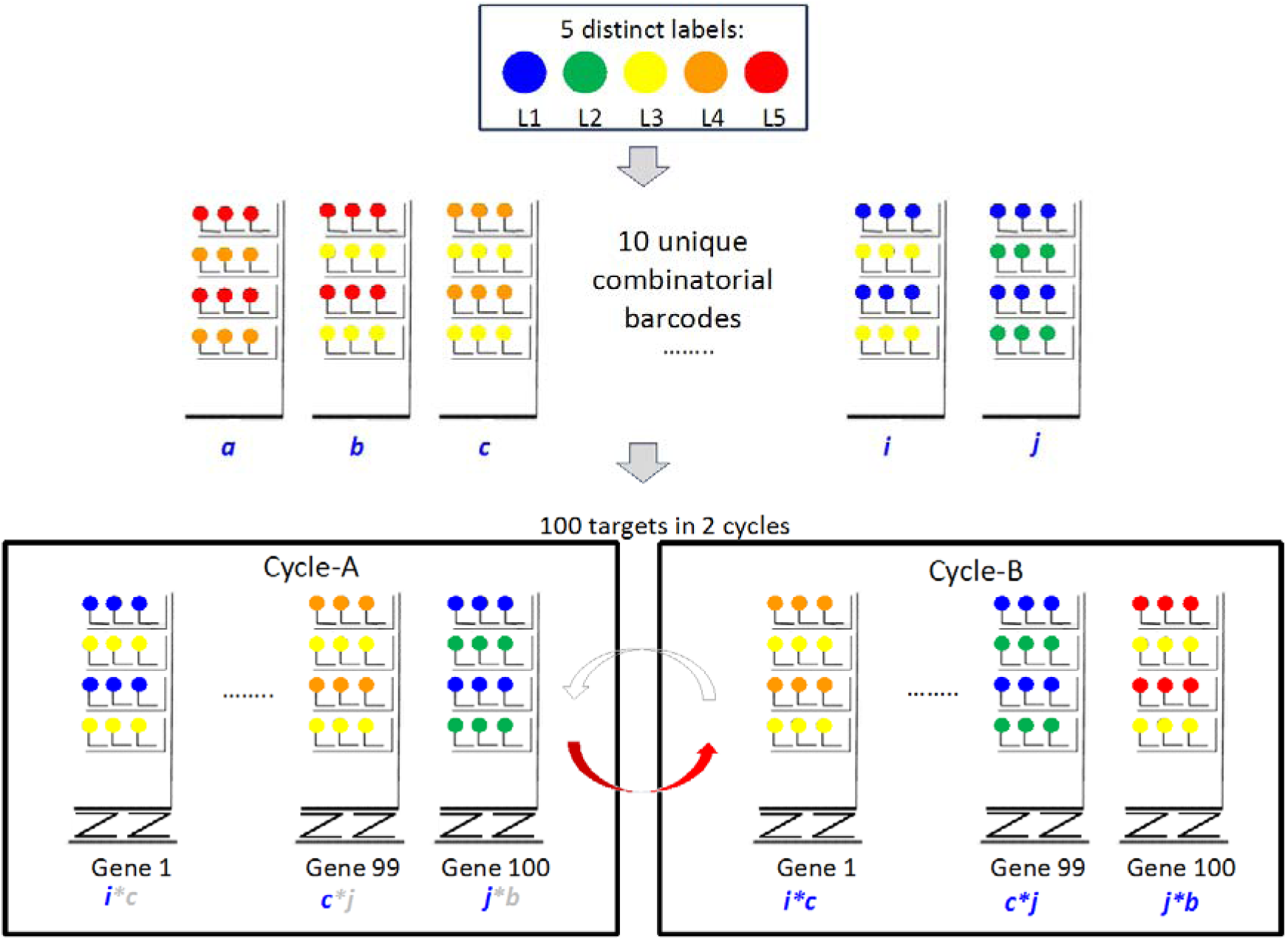
Examples of barcoded detection of target nucleic acids, by generation of two separate independent identifiers of each target nucleic acid using two separate rounds of labeling and detection. The identifiers are incorporated at the label-probe level, using the same region of each target gene in each round. In the first round, each gene is labeled with one type of the signal “trees,” each of which provides signals that form a unique combinatorial code. For example, tree “a” includes only the fourth and fifth label probes, but not the others. Once these signals are detected and recorded to generate the first independent identifier of each target gene, the signals are removed, and the gene is labeled with another type of tree. The second set of signals is then detected and recorded. This procedure can continue with more rounds. The expanded code can then be used to identify the gene.

### Tissue Processing

#### Protease pretreatment for RNA target retrieval

MSIA assay is designed to be compatible with both frozen and FFPE tissue sections. The pretreatment conditions are dependent on the tissue types, tissue processing methods (e.g. frozen; FFPE), and extent of formalin fixation.

### FFPE tissues and autofluorescence reduction

For FFPE tissues, the assay requires a heat-induced target retrieval using RNAscope target retrieval reagent if conducted manually, or Leica Bond ER2 if conducted on Leica Bond system. Following target retrieval, a custom pretreatment reagent developed by Advanced Cell Diagnostics (ACD) was used in the tissue pretreatment step to further untangle the over-crosslinked proteins and expose the mRNA transcripts in the tissue sections. In this study, FFPE mouse brain tissue was first deparaffinized in 100% xylene, dehydrated in 100% ethanol, heat treated with 1X RNAscope target retrieval buffer at 100 °C for 5-10min in a steamer, and then incubated with 300µL of pepsin solution (Sigma-Aldrich) at room temperature for 10min. This pepsin-based pretreatment offers some unique benefits over other conventional proteases from RNAscope pretreatment kits, including reduction in autofluorescence background in FFPE tissues, removal of >90% red blood cells, and removal of non-specific signals, all of which lead to a higher signal-to-noise ratio.

### Fixed-frozen tissues

For fixed-frozen mouse brain sections, the standard RNAscope pretreatment conditions recommended by RNAscope user manual were adopted in the MSIA assay. The fixed-frozen tissues were sectioned at 10uM thickness with cryotome (Leica, Germany), then post fixed in 10% NBF for 15min at RT, followed by a 5-min target retrieval in 1X RNAscope target retrieval buffer, and ACD protease-III digestion for 30min at 40 Celsius.

Slides were incubated with a mixture of 100 MSIA RNA probes for 3hrs at 40°C immediately after the pretreatment.

### ProximityScope assay

To map neuroligin-neurexin interaction at the synaptic cleft, antibodies against Neurexin-3 (Cell Signaling Technology, 48004), Neuroligin-2 (Abcam, Ab315095), and Neuroligin-3 (Novus Biologicals, NBP1-90080) were conjugated with signal-generating oligonucleotides via oYo-Link (AlphaThera, Philadelphia, PA). FFPE mouse brain was pretreated as described in the tissue processing section, stained for 53 RNAs and then incubated with conjugated antibody pairs (Neurexin3-Neuroligin-2, or Neurexin3-Neuroligin3). The oligonucleotide-conjugated antibody pairs were then annealed with 3 tiers of branching DNA oligos (i.e. AMP1, AMP2, and AMP3). The first tier of branching DNA oligos can anneal to antibodies only when both antibodies are in close proximity, thereby ensuring high assay specificity [13]. The third AMP oligo contains a unique binding sequence that is complementary to the corresponding HRP-conjugated oligos, which then activates TSA-conjugated ClariTSA 650 dyes (Tocris, Minneapolis, MN).

### Chromatic Aberration Correction

Accurate assay results rely on precise signal colocalization with subpixel precision. However, chromatic aberration from microscope optics introduces spatial shifts between different fluorophore channels, potentially causing decoding errors. To correct these shifts, we prepared a HeLa cell slide labeled with a single gene barcode (‘11111’) that includes all fluorophores used in the assay (except for the DAPI channel, which does not contribute to color-based colocalization). Since only a single gene is present, all fluorescent signals should ideally colocalize, and any observed displacement represents chromatic aberration.

Because chromatic aberration varies across the field of view (FOV), each FOV was divided into a 5×5 grid (25 sub-FOVs). The channel shifts within each sub-region were measured relative to the first channel (typically 520 nm), and correction values were obtained.

For actual experimental images, the same 5×5 subdivision was applied. Each sub-FOV was corrected using the corresponding reference shifts, and then the corrected sub-FOVs were stitched back together to reconstruct a fully corrected FOV. This corrected image was then used for downstream analysis. Since chromatic aberration is intrinsic to the imaging system, once determined, these correction values remain applicable to subsequent experiments performed on the same setup.

### Stitching and Registration

Since the sample size typically exceeds the microscope’s field of view (FOV), multiple FOV images must be stitched together to generate a complete slide image. Additionally, our assay involves multiple imaging rounds, requiring precise registration of images across rounds to obtain a complete dataset.

For stitching and registration, we largely followed the approach described by Muhlich et al. [14]. Briefly, stitching begins by arranging all FOVs in the DAPI channel according to their recorded stage positions. We then refine these positions by calculating pairwise alignments using phase correlation. To filter out poor alignments, a permutation test is performed by randomly selecting non-adjacent FOVs and determining a threshold to exclude unreliable alignments. A minimum spanning tree is then constructed, using the reciprocal of the correlation values as edge weights. If multiple disconnected trees are formed, the largest tree is used to fit a linear model, which is then applied to assign initial positions for the smaller trees and other disconnected tiles. The tiles within these smaller trees retain their relative positions based on their own measured alignments, ensuring proper placement in the global coordinate system.

For round-to-round registration, the process is more straightforward due to the one-to-one correspondence between images from different rounds. Phase correlation is used to calculate alignment shifts, applying the same threshold derived from the stitching step. Tiles with poor alignment are assigned positions based on a linear model fitted from the successfully aligned tiles.

### RNA Signal Prediction and Decoding Method

The deep-learning framework of ACDnet is based on the feature-pyramid-network equipped with Res50Net architecture that excels at image classification [15]. Inspired by earlier work on designing learning networks that operate on cellular features [16], ACDnet adopts a novel approach by representing dot-like feature prediction outputs as two sets, one for the pixel-wise transform and a second for the inner distance transform.

To address the challenge of lacking ground truth of gene transcript copies per cell, ACDnet adopts a novel approach by representing RNA fluorescent dots as an object with Gaussian shape profile (spatial normal distribution). One of the major benefits is that we can apply the Gaussian distribution function to simulate the complex morphological spatial profiles of RNA fluorescent dots. In addition, by integrating the simulated training data with the artifacts present in the experimentally generated data, we can break the limit of the model’s performance in real data. To train and validate the model, 1024 images were applied for RNA dot training, 256 images were applied for validation and over 128,000 dots were assessed for both training and validation. In essence, we could synthesize a high-quality ground-truth dataset without a tremendous time-consuming and labor-intensive process, to train our deep-learning models.

To achieve higher calling rate without error-correction scheme, we developed Dot-Object-Intensity algorithm and Single-Cell-Zone algorithm to pick up the relatively weak MSIA RNA signals from multiple optical channels (Supplementary Figure. S5). After RNA dot prediction, the identified RNA fluorescent signal dots across multiple imaging cycles that fall into the alignment zone are referenced to a specific binary code by the corresponding cycle order. The alignment was determined by a searching radius threshold with a default value of 4 pixels. Then the resulting binary barcode is compared to the codebook associated with specific RNA marker gene (Supplementary Figure. S6). Due to the efficiency and accuracy of our 5-bits/2-cycles barcoding strategy, no error correction scheme was deemed necessary and thus was not implemented in the decoding process.

### Spatial Mapping of MSIA brain cell types in tissue contexts

For spatial mapping, we take advantage of the scRNA-seq Taxonomic Groups from Allen Brain Cell Atlas [17, 18] and the integrated clustering analysis method from MERFISH [19, 20]. The workflow for spatial mapping of cell types in the ACD-100-plex data is composed of preprocessing (normalization and logarithmic transformation), dimensionality reduction (using principal component analysis, or PCA), neighborhood calculation, Leiden clustering, and cluster matching (using cosine distance).

The key steps of the pipeline are as follows:

1. Dimensionality reduction of ACD-100-plex: ACD-100-plex spatial data, which includes gene expression information at the single-cell level within a spatial context, is subjected to PCA. This transforms high-dimensional gene expression data into a lower-dimensional space, making it easier to visualize and analyze.
2. Clustering and cluster matching: The principal components are then used in a BBKNN (batch balanced k nearest neighbors) analysis to compute a nearest neighbor graph [21]. This nearest-neighbor graph is subsequently used in the Leiden algorithm for clustering. For cluster matching, we use cosine distance between the ACD-100-plex and scRNA-seq data. Cosine distance is a metric used to measure the similarity between two vectors. In this context, it is used to compare the gene expression profiles (represented as vectors) of the ACD-100-plex clusters and scRNA-seq clusters. The smaller the cosine distance, the more similar the two clusters are. The ACD-100-plex cluster is then assigned the taxonomic cell type of the most similar scRNA-seq cluster.

This ACDnet workflow allows for systematic integration of ACD-100-plex spatial data with scRNA-seq data. It enables the transfer of cell type information from scRNA-seq to ACD-100-plex, allowing researchers to spatially map and characterize cell types in tissue contexts.

### Language model for PD-relevant novel gene discovery

The language model used in this study is an embedding model based on a Skip-gram architecture, which has been shown to outperform bag-of-words approaches in capturing contextual relationships [22]. The embedding dimensionality was set to 100, providing a balance between model complexity and computational efficiency. To emphasize semantic relationships between terms, a relatively large context window size of 6 was selected. Negative sampling was employed to improve both computational efficiency and model performance, consistent with previous findings [22, 23], with the number of negative samples set to 5.

The model was trained with a learning rate of 0.001, utilizing the Adam optimizer to ensure stable convergence and effective regularization. A batch size of 8192 was used during training to optimize computational performance while maintaining robust gradient estimation. The training corpus was compiled from full-text articles and abstracts retrieved from PubMed and PMC, using the search term “Parkinson’s disease (MeSH) AND gene” to capture relevant literature published between 2014 and 2023.

## Results

### MSIA achieved highly specific in-situ mRNA detection and enables simultaneous transcriptome profiling at high-throughput

To determine the fidelity of MSIA assay in comparison to established commercial methods, we subjected one coronal section collected from a 1-month-old frozen mouse brain to 100-gene MSIA and compared the in situ profiling results to a publicly available *in situ* spatial transcriptome datasets generated using Molecular Cartography (Resolve Biosciences, Germany) [24], as well as to a 234-genes spatial transcriptomic dataset of an age-matched mouse brain section generated by the 10x Genomics Xenium assay [25]. This head-to-head direct comparison was executed by our ACDnet pipeline, and the results are presented in **Figure 2**. Both Xenium and MSIA technologies revealed comparable transcript copies/cell for low and medium expression genes (e.g. genes <3 copies/cell in Xenium. **Figure.2 A,C**). For higher expression genes (e.g. genes over 3 copies/cell in Xenium), MSIA assay detected fewer transcript copies/cell than the Xenium (**Figure.2 A,C**). To validate the copy numbers of these high-discrepancy genes in mouse brain, we conducted RNAscope HiPlex assay, an established validation method [26], using RNAscope probes to target the same gene transcript species on consecutive sections collected from the same 1-month-old fixed-frozen mouse brain.

**Figure 2.**
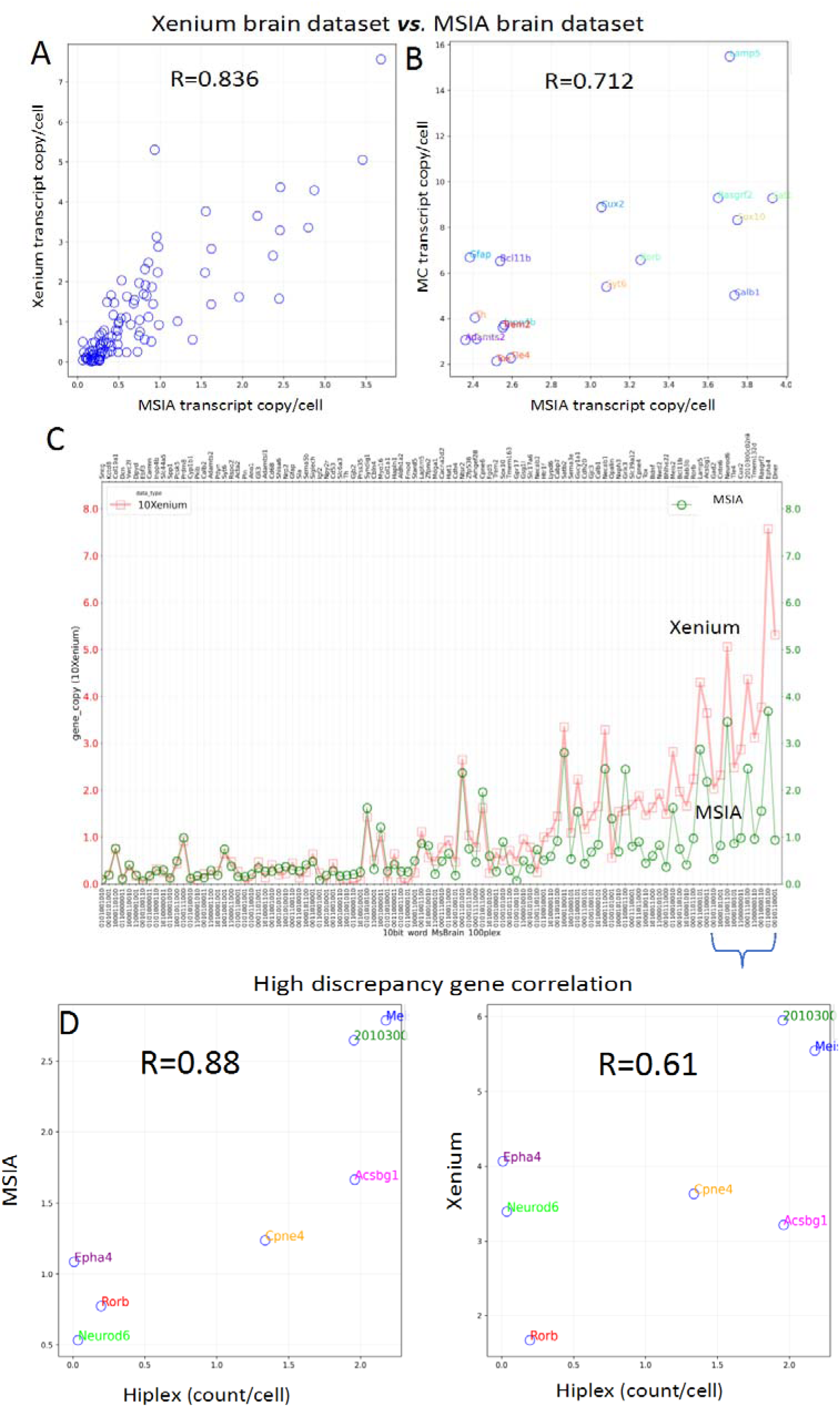
Comparative analyses of MSIA spatial transcriptomic profiling performance. A-B: Correlation between the Xenium 1-month-old mouse brain dataset and MSIA 1-month-old mouse brain dataset in gene transcript copies/cell. The two datasets showed comparable transcript copies/cell for lower and median expression targets despite larger discrepancies in the last 10 genes (B, bracketed genes). These genes showed higher expression levels in Xenium dataset but lower copies/cell in either MSIA assay or orthogonal HiPlex assay (C and D).

Quantification of the RNAscope HiPlex signals revealed the HiPlex gene copy numbers (i.e. average dot count/cell) exhibited a stronger linear correlation with the orthogonal MSIA data (R=0.88; **Figure.2D**), as compared with Xenium gene copy numbers (R=0.61; **Figure.2D**). It should be noted that the 10x dataset was created in 2022 with Xenium chemistry v1; 1.0.0.3 instrument software; Xenium-1.0.2 analysis software, and a software update was reported to correct the overestimation of high expressors in Xenium since performing these MSIA experiments. Consistently, A comparison between the Molecular Cartography brain dataset and our MSIA dataset showed that the two datasets share seventeen genes in common, with an R value of 0.71 (**Figure 2B**).

Overall, MSIA yields reliable gene expression measurements for low, medium and high expressors in this set of experiments. In some experiments, a tissue clearing procedure was deployed first before MSIA assay, to reduce high autofluorescence from over-crosslinked protein structures as well as blood cells (Supplementary Figure. S2).

### MSIA yields accurate *in situ* cell type characterization and mapping in mouse brain sections

With only 100 differential neuronal and non-neuronal gene markers, the semi-automated assay accurately identified the major brain cell classes in different age groups (28∼30 cell clusters/brain; **Figure.3 D-F**), unveiled a spatial cell atlas of the whole mouse brain (**Figure.3 A-C**), depicting the highly organized neuron lamination and their orchestrated interactions with glial, vascular leptomeningeal cells (VLMCs) immune, and other non-neuronal cells (**Figure.3 J-L**). The spatial affinities between neuron and nonneuronal cells are consistent with previous findings [17, 20, 27] and demonstrated an active remodeling of neural connectivity and functions of glial cells in the life span of the mice. To confirm the spatial locations of the brain cell classes, we also performed immunofluorescent staining against canonical cell type markers in the third cycle of MSIA following the RNA detection cycles (**Figure. 3M** and **Supplementary Figure. S3**). NeuN-immunopositive neurons showed a spatially dense distribution comparable with that of MSIA neuron cells which are transcriptomically defined through aligning their transcriptional signatures to a single-cell RNA-seq brain cell atlas (**Figure.3 M;** see Methods for details.).

**Figure 3.**
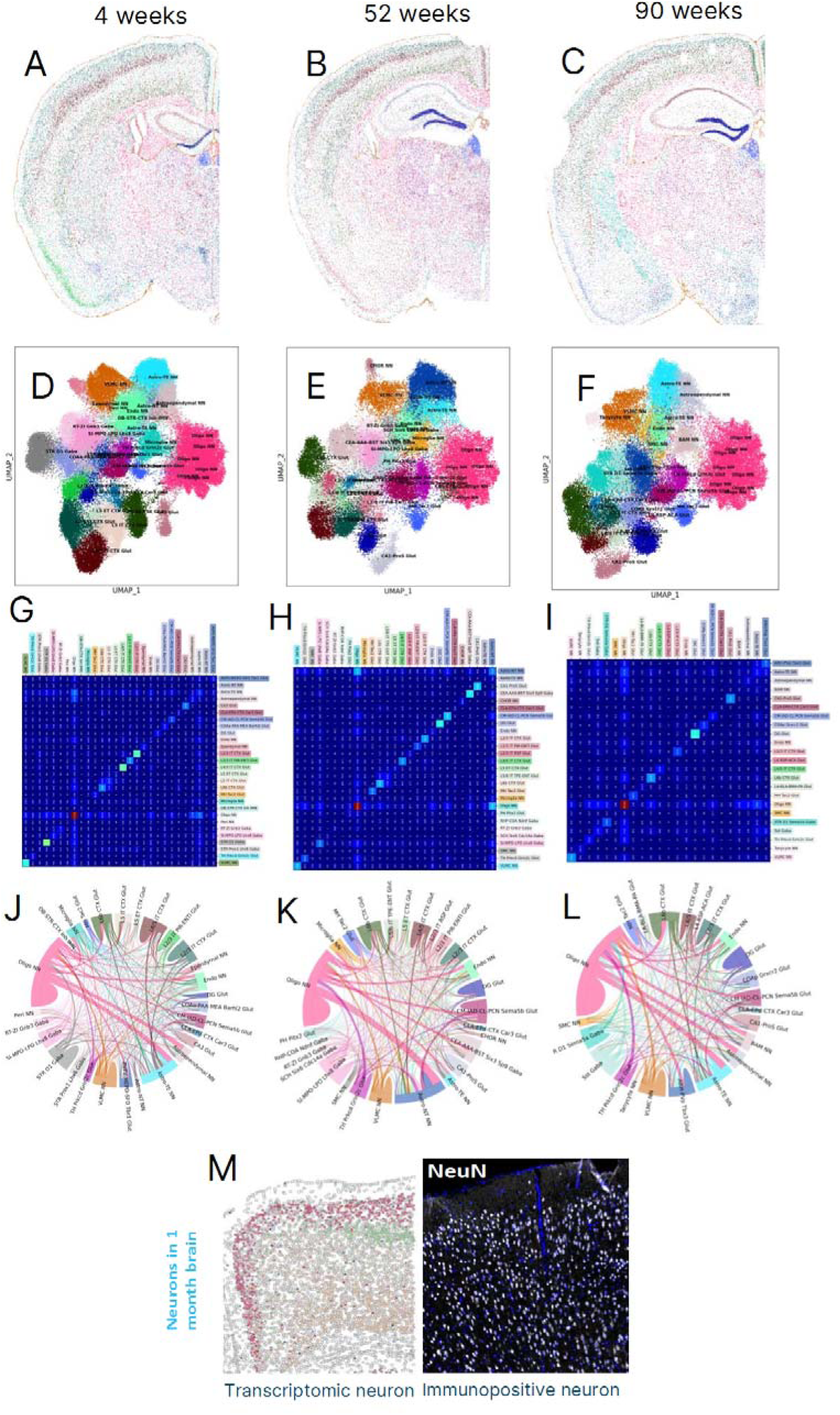
Spatial transcriptomics analysis of mouse brains at different life spans. (**A-C**) spatial atlas of MSIA cell classes of mouse brains at the age of 4, 52, 90 weeks post-natal, defined molecularly by our MSIA 100 RNA spatial transcriptomics data. (**D-F**) UMAP projection of MSIA cells annotated using unsupervised labels derived from sc-RNAseq dataset identifies 23-28 clusters in the aging brains. (**G-I**) Heatmap showing the correlation scores of different pairs of neighboring brain cell classes. (**M**) comparison of the spatial locations of MSIA neurons and the immunopositive neurons in the same brain section (MOp).

Furthermore, the spatial distribution of transcriptomic microglia/macrophage cells in a brain cortex MOp region was validated with corresponding protein markers CD68 and Iba1. Likewise, CD68+ and Iba1+ cells exhibited a spatial distribution comparable with the corresponding MSIA cell population (**Supplementary Figure. S4.A**). In contrast, GFAP immunopositive astrocytes showed a highly localized distribution (**Supplementary Figure. S4.B; right panel**), while the spatial distribution of astrocyte defined by single-cell spatial transcriptomics showed a broader distribution across different cortical layers (**Supplementary Figure. S4.B; left panel**). These data may suggest the complexity of epigenetic regulations and cell state change in response to environmental stimuli and aging process.

### MSIA in-depth transcriptome profiling revealed novel molecular mechanisms in MPTP-induced murine brain model of Parkinson’s disease

MSIA yielded results that were consistent with previous findings in mouse brain aging and provided supporting evidence to deploy the technology for PD translational research. We chose to further understand the molecular mechanism of PD in mouse models because of the abundance of available data that can be used to further validate our method and used a language model trained on PD literature to identify PD related differentially expressed genes that can be validated on human samples. We generated a PD mouse model by acute 1-methyl-4-phenyl-1,2,3,6-tetrahydropyridine (**MPTP**) injections. MPTP mouse model is a well-established PD model that is marked by large dopaminergic neuronal death in substantia nigra and dopamine depletion of the striatum. Both C57BL/6 mice were treated with MPTP or vehicle via repeated intraperitoneal injections on day 0 and day 1. Mouse whole brain was harvested on Day 8 and processed immediately into FFPE samples. Body weight of the MPTP-treated mice decreased notably in day 1 and day 2 due to the MPTP dosing and started to increase starting day 3. No significant body weight differences were observed between MPTP-and vehicle-treated mice throughout the whole study period. To assess the motor skills of these mice, fine motor kinematic gait analysis was performed on day 7. Typical

MPTP-induced kinematic phenotypes were observed, including increased hip and knee angles and tail tip height, increased homologous and diagonal gait, altered forelimb trajectory profiles, and increased vertical hip movements. The Gait Overall Score of the MPTP-treated mice is significantly higher than that of the vehicle treated mice (**Supplementary Figure. S1**). Overall, these data suggested that we successfully induced Parkinsonism in the MPTP injected mice.

We then curated a panel of 53 DA neuron-related gene markers to characterize DA neuron degeneration in MPTP mouse brains. These gene markers were meticulously selected from recent single-cell RNA sequencing datasets in normal and PD human brain samples and WT and MPTP mouse brain samples [27–32] in effort to sufficiently depict the disease status and reduce spatial crowding. Using this 53-plex MSIA panel, we transcriptionally profiled the substantia nigra (SN) region in the vehicle and MPTP-treated mouse brains. The assay revealed extensive reductions in various brain cell types and subtypes, including DA neurons, pericytes, resident macrophages, and Oligodendrocyte progenitor cell (OPC), while astrocyte, oligodendrocyte, and MB-Glu neurons appeared to be less affected by the MPTP treatment (**Figure.4 B-C**).

Consistent with previous results, significant DA neuron loss was observed in substantia nigra region of the MPTP-treated mouse brain (**Figure.4 D-E**). Among the 53 panel genes, our assay found the mRNA expressions of 17 genes were evidently affected by MPTP treatment while the other genes remained in comparable expression levels in WT and MPTP brain SN cells. (**Figure. 4 F**). Specifically, the gene expressions of 2 DA neuron markers (Slc6a3 and Anxa1) were downregulated by the MPTP treatment (**Table.3 DA neuron**). Genes that are biologically associated with other neuronal and non-neuronal cell types and genes closely associated with PD progression were also differentially expressed in the MPTP treated mouse brains (**Table.3**) [27–34]

**Figure 4:**
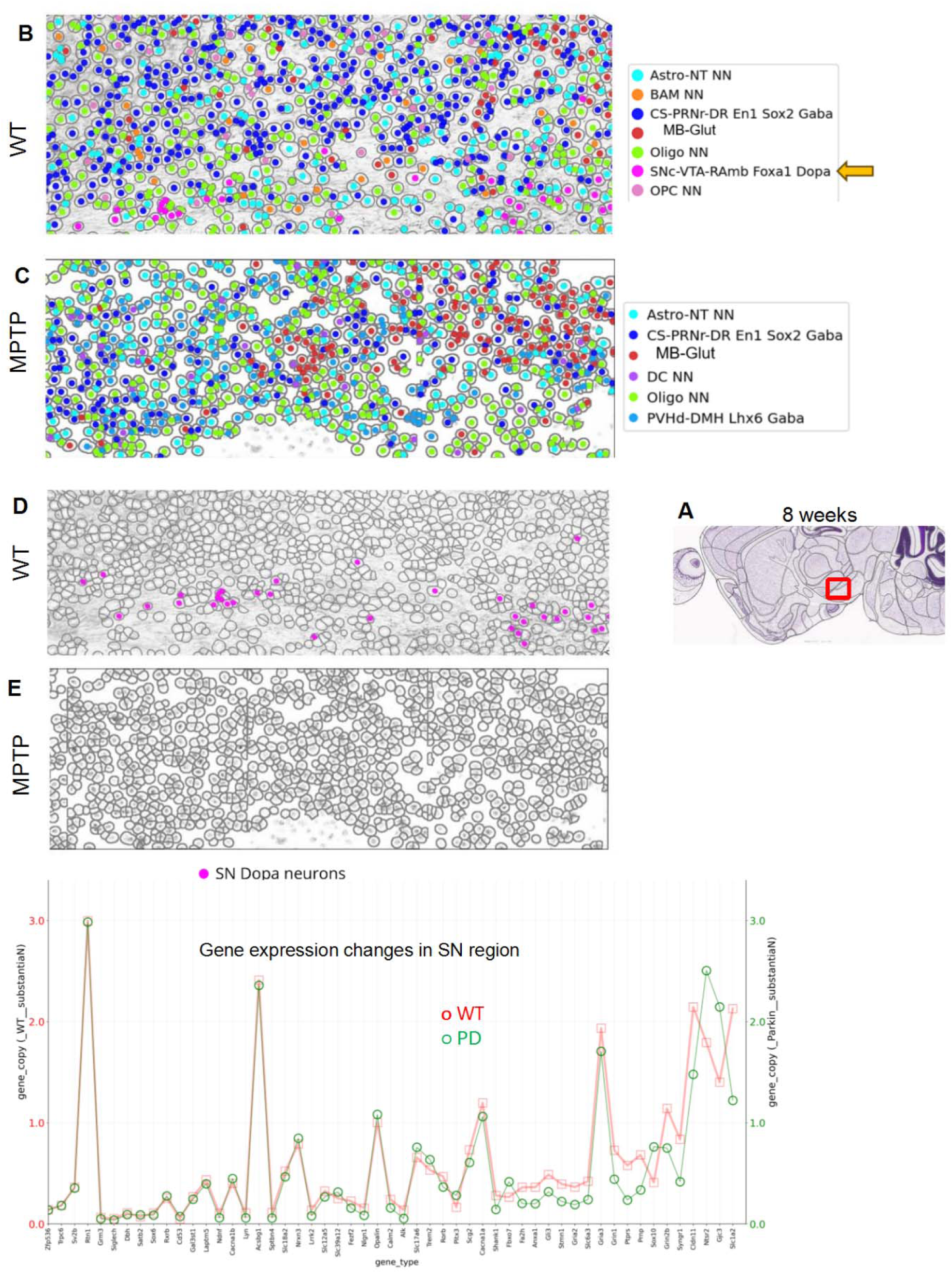
MSIA assay revealed transcriptional and cell type changes in MPTP-treated mouse brain as compared with WT brain. (A) the SN region in sagittally sectioned mouse FFPE brain slides (WT and PD, 5μM thick) were selected for MSIA cell type analysis. (B-C) spatial distribution of different MSIA transcriptomic cell classes in SN region. (D-E) spatial representation of the SN DA neurons in WT midbrain vs. MPTP-treated midbrain. (F) changes in mRNA expressions in mouse SN following MPTP treatment (unit: average transcript copies/cell)

### Spatially resolved neurexin-neuroligin protein ProximityScope™ assay (PSA) map revealed a compromised synaptic interaction among DA neurons and other neuronal types in SN regions of mouse PD brains

Neurexins (NRXs) and Neuroligins (NLGNs) are key protein players in synapse formation and maintenance. NRXs, as presynaptic cell adhesion molecules, are involved in the synaptic specification and differentiation of excitatory and inhibitory synapses. NLGNs, as postsynaptic proteins, regulate synaptogenesis and maintain synaptic stability. During development, neuroligins contribute to the fine-tuning of neural circuits. Mutations and deletions in the neurexin or the neuroligin genes have been linked to psychiatric disorders like autism, epilepsy, schizophrenia, and neurodegenerative disorders. The interaction between neurexins and neuroligins in the synaptic cleft is responsible for effective synaptic transmission and plasticity in learning and memory. To further demonstrate the multi-omics capacity of MSIA and provide insights on the synaptic interaction landscape in response to MPTP treatment, we conducted a 3-cycles MSIA assay in the WT and MPTP mouse brains. Briefly, FFPE sagittal brain sections first underwent two cycles of RNA optical barcoding and signal removal, followed by a third cycle of neurexin-neuroligin protein interaction staining was conducted on the same slides to visualize the physical interaction between the NRX/NGLN adhesion complexes of the whole brain. Our study showed that, the mouse PD brains, induced by MPTP intraperitoneal injection (80mg/kg), exhibited an extensive disruption of synaptic NRXN3-NLGN2 interactions compared to age-matched, vehicle control brains in HCF (**Figure. 5 A-B**), midbrain (**Figure. 5 C-D),** and striatum (**Figure. 5 E-F**) (Arrowhead= NRXN3-NLGN2 protein complex in synapses). Similarly, PD mouse brain also exhibited significantly weaker NRXN3/NGLN3 interactions compared with the age-matched, wild type brains, as indicated by the MSIA protein staining cycle specifically targeting the NRXN3/NGLN3 interaction.

**Figure. 5:**
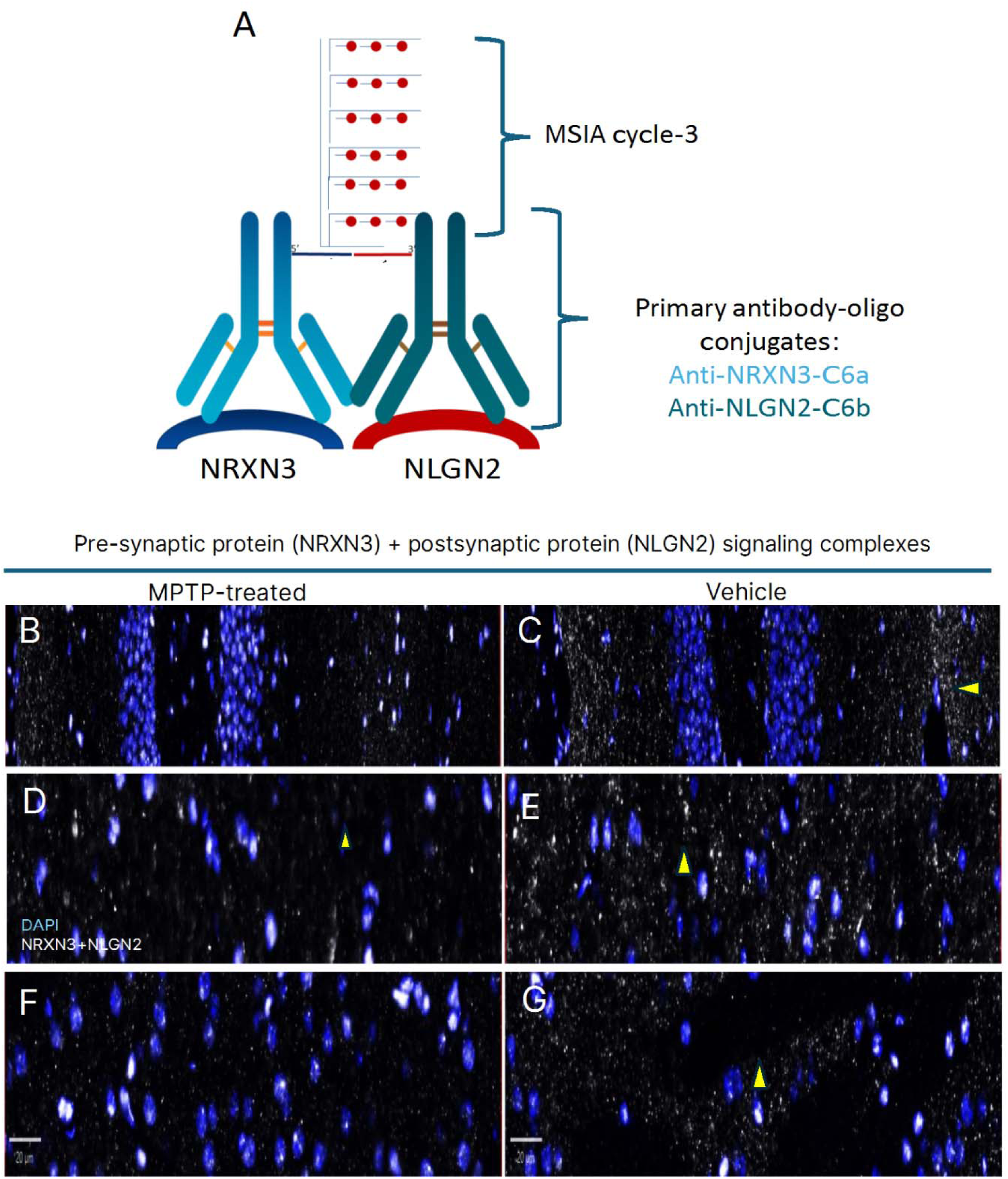
MSIA combined with a proprietary protein-protein interaction assay visualized the spatial distribution of neuron synaptic signaling complexes in mouse brains at cellular resolution. (A) in the third cycle, a ProximityScope™ assay was performed on the cleaved slide using oligo-conjugated primary antibodies against Neurexin-3 and neuroligin-2. The additional protein-protein interaction cycle provided an *in situ* visualization of neuron synaptic structures, organization, and the extent of neuron-neuron interactions in the whole brain section (B-D). Mouse model of Parkinson’s disease induced by MPTP I.P injection (80mg/kg) showed extensive disruption of synaptic NRXN3-NLGN2 interactions in HCF (B-C), midbrain (D-E), and striatum (F-G) compared to age-matched, vehicle control brains (*Arrowhead= NRXN3-NLGN2 protein complex in synapses*).

### Integrative Gene Selection Strategy Combining Language Model Predictions and Expression Databases for Differential Analysis in WT and PD Mouse Brains

From the initial pool of 53 genes tested using MSIA, we identified 11 genes that exhibited observable differences in expression between WT and PD mouse brains. These 11 genes were further filtered based on their frequency in the training data of the embedding model, applying a threshold of 50, which reduced the list to 6 genes (Fbxo7, Grin2b, Prnp, Slc1a2, Slc6a3, and Slc18a2). The threshold of 50 was chosen to ensure that the embedding model had sufficient exposure to each gene during training, which is critical for generating reliable representations. A higher threshold would risk narrowing the candidate list excessively, while a lower threshold might include genes with less robust embeddings due to insufficient contextual information. Using the embedding model, we predicted the top 100 genes most similar to each of these 6 genes. For each of these six genes, we also identified another set of the top 100 closest genes based on their expression patterns [35]. Genes that appeared in both lists were selected, and across all six genes, this process resulted in a total of 25 candidate genes for further analysis.

These 25 candidates were then cross-referenced with a database[36] that reports gene expression levels in both WT and PD mouse brains. We selected genes that showed consistent expression changes across multiple samples in the database, narrowing the list to 9 genes. Finally, we included the control gene Polr2a, which exhibits stable expression between PD and WT, bringing the total to 10 genes. These 10 genes were incorporated into a panel for experimental validation using the FISH assay.

We analyzed the expression of 10 selected genes, including a control gene (Polr2a), in WT and PD mouse brain samples using the RNAscope HiPlex v2 assay (Bio-Techne). Among these, four genes (Drd2, Kcnj6, Sv2c and Polr2a) produced robust signals (**Figure. 6**), and their expression levels were quantified at the single-cell level in the substantia nigra (SN) using Halo v3.5 (Indica Labs). For the remaining six genes, we employed the RNAscope Multiplex Fluorescent V2 assay (Bio-Techne) optimized for low-expression targets, followed by Halo-based quantification (**Figure. 7**). Of the 10 genes, Chrna3 and Usp8 exhibited expression levels too low for reliable quantification in the SN region. For the remaining eight genes, six showed (Kcnj6, Drd2, Polr2a, Ankrd34b, Sv2c and Elavl2) expression trends consistent with what is reported in the literature[36], supporting the effectiveness of our methods presented here. The other two genes, Srebf1[37] and Mbnl2[38], have not been reported to show changes in gene expression in an MPTP mouse model and any marginal changes that we observe in our experiments have to be confirmed in additional animals.

**Figure 6.**
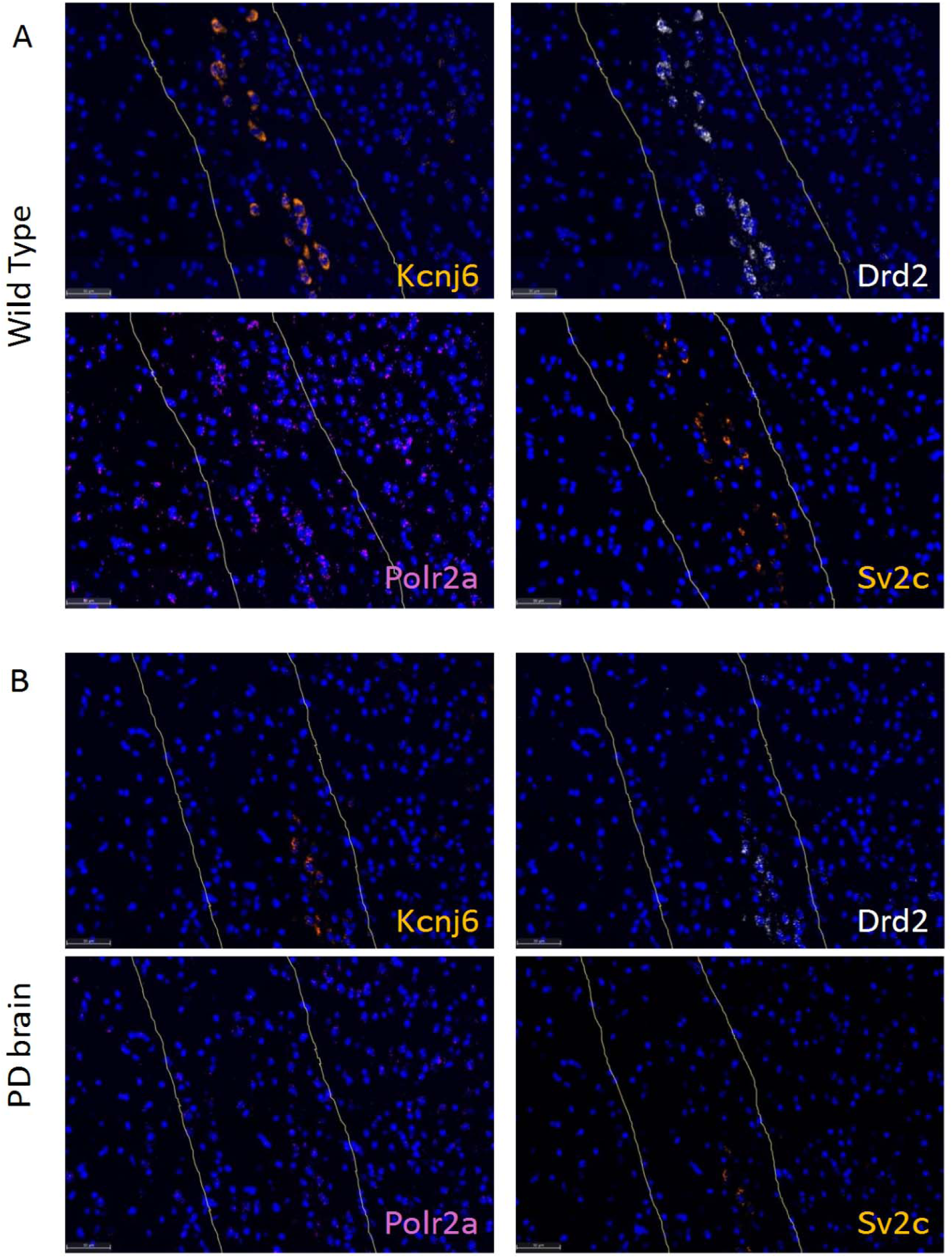
The expression of four genes exhibiting a strong signal in the RNAscope HiPlex v2 assay. (A) Visualization of *Kcnj6*, *Drd2*, *Polr2a*, and *Drd2* in a brain section from a wild-type mouse, with the substantia nigra region highlighted by the yellow outline. (B) Visualization of *Kcnj6*, *Drd2*, *Polr2a*, and *Drd2* in a brain section from a Parkinson’s disease model mouse.

**Figure 7.**
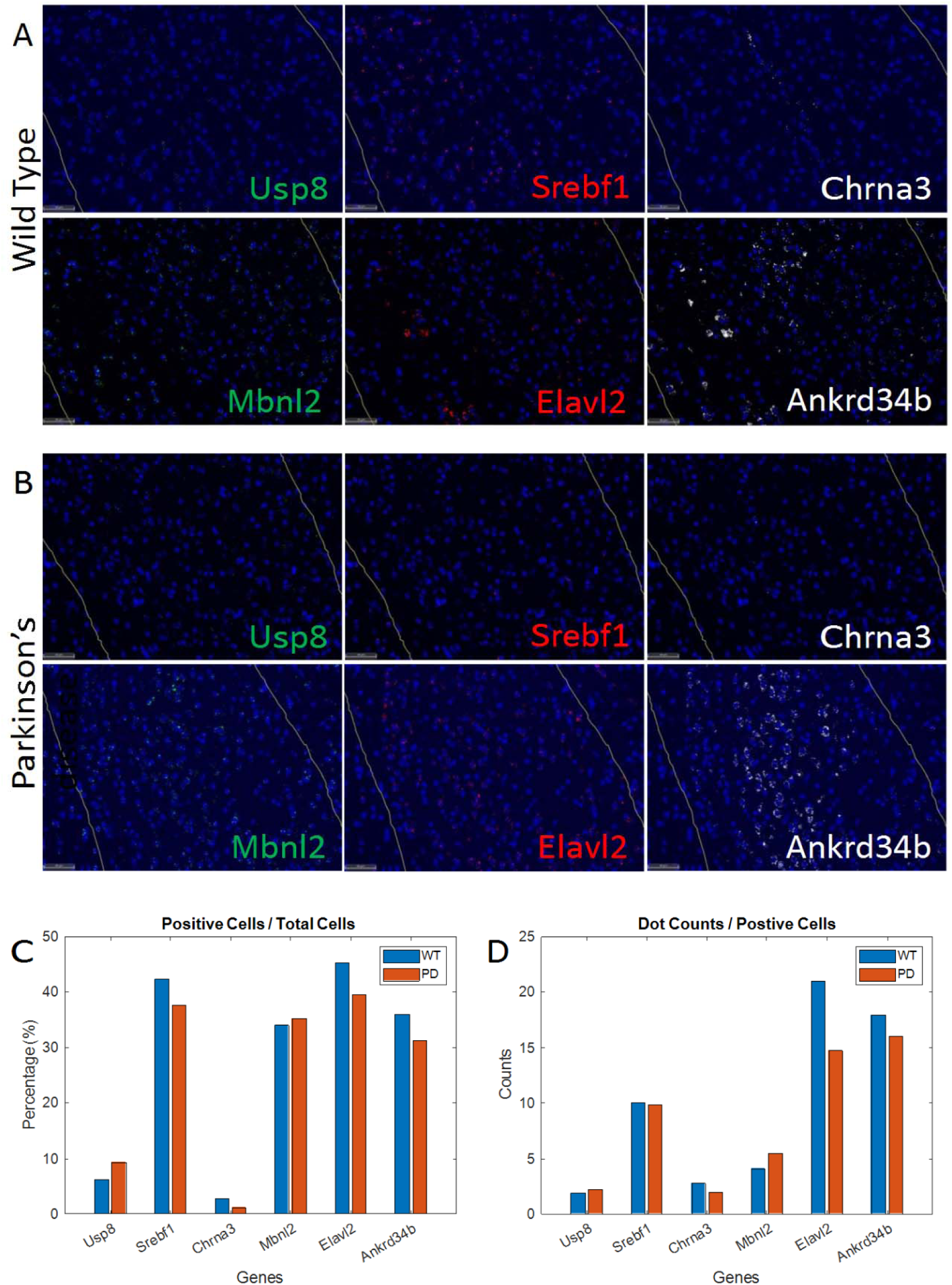
The remaining six genes were detected by RNAscope Multiomics LS assay. (A) Visualization of *Usp8*, *Srebf1*, *Chrna3*, *Mbnl2*, *Elavl2*, and *Ankrd34b* in a brain section from a wild-type mouse, with the substantia nigra (SN) region outlined in yellow. (B) Visualization of the same genes (*Usp8*, *Srebf1*, *Chrna3*, *Mbnl2*, *Elavl2*, and *Ankrd34b*) in a brain section from a Parkinson’s disease model mouse. (C) Bar graph comparing the percentage of gene-positive cells relative to the total cell numbers in the SN region between wild-type (WT) and Parkinson’s disease (PD) mice. (D) Bar graph showing the dot count per positive cell for each gene in the SN region, comparing WT and PD cases.

## Discussion

In this report, we demonstrate the development of MSIA, a novel in situ hybridization method with 5-color optical barcoding to enable the detection of 100s of expressed genes. We tested this method initially on fresh frozen mouse brain samples of different ages and reproduced age-dependent gene expression changes that were previously reported. This method has several advantages – a manual workflow composed of just two stain-image cycles that obviates the need to purchase expensive instrumentation, custom filter sets that can be fitted to any conventional fluorescent microscope, modified RNAscope chemistry that results in high demonstrated sensitivity especially for low expressors and an in-house developed deep learning-based data analysis pipeline for dot detection trained on hundreds of RNAscope images. We then compared gene expression changes in 53 genes between wild type and a Parkinson’s disease mouse model. We also performed a third round of stain-image cycle where we stained for cell markers NeuN, GFAP, CD68 and Iba1 showcasing the compatibility of multiomic workflows. In addition, we used the newly developed protein-proximity assay to assess the integrity of synapses using NRXN3-NLGN2 proximity and, discovered severe loss of synapses in the substantia nigra region of PD mouse brains, reproducing previous studies [39].

Software and image analysis methods were designed using a combination of publicly available tools and proprietary RNAscope images that resulted in a reliable RNA dot counting algorithm. The fidelity of the algorithm was assessed by validating the findings from MSIA on the established RNAscope HiPlex method as well as reported expression levels in published scRNAseq data of mouse brains.

To translate findings to human samples and discover biomarkers for Parkinson’s disease, we identified 11 genes that exhibited significant differences in expression between WT and PD mouse brains from the initial pool of 53 genes tested using MSIA. We then developed a language model trained on PD literature. The 11 genes further filtered out based on their frequency in the training data of the language model to generate a list of 6 genes of high significance (Fbxo7, Grin2b, Prnp, Slc1a2, Slc6a3 and Slc18a2). Each of these genes is highly relevant to PD as they regulate key processes like protein degradation (Fbxo7) [40], excitotoxicity (Grin2b, Slc1a2) [41–44], dopaminergic neurotransmission (Slc6a3, Slc18a2) [45, 46], and protein aggregation (Prnp) [47]. We then identified the top 100 genes that were similar to each of these 6 genes from the language model as well as the top 100 differentially expressed genes between WT and PD mouse models from a gene expression database, compared the two lists to generate 9 high confidence genes that were further studied in human normal and PD samples.

Finally, we believe MSIA easily lends itself to a throughput of over 1000 genes by the use of additional stain/image cycles in combination with more colors and a different

Hamming weight, potentially with an error-correction mechanism included. Commercially available HiPlex methods use three stain-image cycles, and within ACD we have extended the number of cycles to eight using HiPlex Up reagent (data not shown). Therefore, addition of one or two additional stain-image cycles to the MSIA workflow has the potential to quantify over 1000 genes. Nevertheless, the current effort is limited to a maximum of ∼100 genes and for the current effort, we have deployed the high sensitivity and specificity of RNAscope chemistry to quantify predominantly low and medium expressors, complementing the strengths of screening technologies like the Visium HD, Xenium and CosMx. Like other spatial imaging technologies, MSIA also requires a predetermined panel of genes (with 10s of barcodes reserved for custom panels that can be added to the predetermined list of genes). The findings from this study would have to be reproduced in more mouse and human samples to add to the substantial body of PD literature. Nonetheless, the availability of a manual and semi-automated MSIA workflows should add to the arsenal of technologies available to interrogate the complex spatial neighborhood of normal and diseased tissues.

## Author Contributions

CZ, AL, JZ, CWC, YW and MS wrote the manuscript, CZ, JZ, YW, CWC, LCW and MS conceived and configured the product concept. CZ, JZ YW and PW performed the experiments. GAK, SD, LCW, MS conceived, designed, performed and launched ProximityScope to detect protein proximity, MY, YY, SC, LD and CZ designed RNA probes used in the experiment. AL, JZ and CWC designed deep learning algorithms, language models and performed data analysis.

## Abbreviations

NBF: neutral buffered formalin
PFA: paraformaldehyde
FFPE: formalin fixed paraffin embedded
PCA: principal component analysis
PD: Parkinson’s disease
MPTP1: methyl-4-phenyl-1,2,3,6-tetrahydropyridine.

## References

1. Liu, X., et al., Spatial multi-omics: deciphering technological landscape of integration of multi-omics and its applications. J Hematol Oncol, 2024. 17(1): p. 72.

2. Morris, H.R., et al., The pathogenesis of Parkinson’s disease. Lancet, 2024. 403(10423): p. 293–304.

3. Ben-Shlomo, Y., et al., The epidemiology of Parkinson’s disease. Lancet, 2024. 403(10423): p. 283–292.

4. Luo, Y., et al., Global, regional, national epidemiology and trends of Parkinson’s disease from 1990 to 2021: findings from the Global Burden of Disease Study 2021. Front Aging Neurosci, 2024. 16: p. 1498756.

5. Cappelletti, C., et al., Transcriptomic profiling of Parkinson’s disease brains reveals disease stage specific gene expression changes. Acta Neuropathologica, 2023. 146(2): p. 227–244.

6. Dikshit, A., et al., Spatial multiplex profiling of immune cell markers in FFPE tumor tissues using the RNAscope™ HiPlex v2 in situ hybridization assay. Cancer Research, 2022. 82(12_Supplement): p. 3865–3865.

7. Rademacher, A., et al., Comparison of spatial transcriptomics technologies using tumor cryosections. Genome Biology, 2025. 26(1): p. 176.

8. Camacho, C., et al., ElasticBLAST: accelerating sequence search via cloud computing. BMC Bioinformatics, 2023. 24(1): p. 117.

9. Advanced-Cell-Diagnostics. RNAscope™ HiPlex12 Reagent Kit (488, 550, 650, 750) v2 Standard Assay with Sample Preparation and Pretreatment. 2022 02/04; Rev B:[Available from: https://resources.bio-techne.com/products/documents/manual/mnl_324440_rnascope-intro-pack-for-hiplex12-reagents-kit-488-550-650-750-mm-v2_20250123033707338.pdf.

10. Janesick, A., et al., High resolution mapping of the tumor microenvironment using integrated single-cell, spatial and in situ analysis. Nature Communications, 2023. 14(1): p. 8353.

11. Moffitt, J.R. and X. Zhuang, RNA Imaging with Multiplexed Error-Robust Fluorescence In Situ Hybridization (MERFISH). Methods Enzymol, 2016. 572: p. 1–49.

12. Advanced-Cell-Diagnostics. Multiplex Assays Using the HiPlexUp Reagent. 2021 10/06; Rev A:[Available from: https://acdbio.com/sites/default/files/QG%20324190revB%20HiPlexUp%20Quick%20Guide_0.pdf.

13. Advanced-Cell-Diagnostics. RNAscope™ Multiomic LS Detection Kit User Manual. 2026 01/08; Rev G:[Available from: https://resources.bio-techne.com/products/documents/manual/mnl_323175_rnascope-multiomic-core-c1-c6-channel-kit_20260112201535038.pdf.

14. Muhlich, J.L., et al., Stitching and registering highly multiplexed whole-slide images of tissues and tumors using ASHLAR. Bioinformatics, 2022. 38(19): p. 4613–4621.

15. Lin, T.-Y., et al. Feature pyramid networks for object detection. in Proceedings of the IEEE conference on computer vision and pattern recognition. 2017.

16. Greenwald, N.F., et al., Whole-cell segmentation of tissue images with human-level performance using large-scale data annotation and deep learning. Nature biotechnology, 2022. 40(4): p. 555–565.

17. Yao, Z., et al., A high-resolution transcriptomic and spatial atlas of cell types in the whole mouse brain. Nature, 2023. 624(7991): p. 317–332.

18. Yao, Z., et al., A taxonomy of transcriptomic cell types across the isocortex and hippocampal formation. Cell, 2021. 184(12): p. 3222–3241 e26.

19. Allen, W.E., et al., Molecular and spatial signatures of mouse brain aging at single-cell resolution. Cell, 2023. 186(1): p. 194–208.e18.

20. Zhang, M., et al., Molecularly defined and spatially resolved cell atlas of the whole mouse brain. Nature, 2023. 624(7991): p. 343–354.

21. Polanski, K., et al., BBKNN: fast batch alignment of single cell transcriptomes. Bioinformatics, 2020. 36(3): p. 964–965.

22. Tshitoyan, V., et al., Unsupervised word embeddings capture latent knowledge from materials science literature. Nature, 2019. 571(7763): p. 95–98.

23. Mikolov, T., et al., Distributed representations of words and phrases and their compositionality. Advances in neural information processing systems, 2013. 26.

24. Jin, K., et al., Cell-type specific molecular signatures of aging revealed in a brain-wide transcriptomic cell-type atlas. bioRxiv, 2023: p. 2023.07.26.550355.

25. Genomics, X., Fresh Frozen Mouse Brain for Xenium Explorer Demo. 2022.

26. He, S., et al., High-plex imaging of RNA and proteins at subcellular resolution in fixed tissue by spatial molecular imaging. Nat Biotechnol, 2022. 40(12): p. 1794–1806.

27. Hook, P.W., et al., Single-Cell RNA-Seq of Mouse Dopaminergic Neurons Informs Candidate Gene Selection for Sporadic Parkinson Disease. Am J Hum Genet, 2018. 102(3): p. 427–446.

28. Kamath, T., et al., Single-cell genomic profiling of human dopamine neurons identifies a population that selectively degenerates in Parkinson’s disease. Nat Neurosci, 2022. 25(5): p. 588–595.

29. Guo, Y., et al., Defining Specific Cell States of MPTP-Induced Parkinson’s Disease by Single-Nucleus RNA Sequencing. Int J Mol Sci, 2022. 23(18).

30. Zhong, J., et al., Single-cell brain atlas of Parkinson’s disease mouse model. J Genet Genomics, 2021. 48(4): p. 277–288.

31. Tiklova, K., et al., Single-cell RNA sequencing reveals midbrain dopamine neuron diversity emerging during mouse brain development. Nat Commun, 2019. 10(1): p. 581.

32. Poulin, J.F., et al., Defining midbrain dopaminergic neuron diversity by single-cell gene expression profiling. Cell Rep, 2014. 9(3): p. 930–43.

33. Yao, X., et al., Differential gene expression and immune profiling in Parkinson’s disease: unveiling potential candidate biomarkers. BMC Neurol, 2025. 25(1): p. 354.

34. Noda, S., et al., Aging-related motor function and dopaminergic neuronal loss in C57BL/6 mice. Mol Brain, 2020. 13(1): p. 46.

35. Zhang, Y., et al., An RNA-sequencing transcriptome and splicing database of glia, neurons, and vascular cells of the cerebral cortex. Journal of neuroscience, 2014. 34(36): p. 11929–11947.

36. Huang, Q., et al., Cell type-and region-specific translatomes in an MPTP mouse model of Parkinson’s disease. Neurobiology of Disease, 2023. 180: p. 106105.

37. Ivatt, R.M., et al., Genome-wide RNAi screen identifies the Parkinson disease GWAS risk locus SREBF1 as a regulator of mitophagy. Proceedings of the National Academy of Sciences, 2014. 111(23): p. 8494–8499.

38. Charizanis, K., et al., Muscleblind-like 2-mediated alternative splicing in the developing brain and dysregulation in myotonic dystrophy. Neuron, 2012. 75(3): p. 437–450.

39. Cuttler, K., et al., Emerging evidence implicating a role for neurexins in neurodegenerative and neuropsychiatric disorders. Open Biol, 2021. 11(10): p. 210091.

40. Joseph, S., J.B. Schulz, and J. Stegmüller, Mechanistic contributions of FBXO7 to Parkinson disease. Journal of neurochemistry, 2018. 144(2): p. 118–127.

41. Cui, C., et al., Association between GRIN2B polymorphism and Parkinson’s disease risk, age at onset, and progression in Southern China. Frontiers in Neurology, 2024. 15: p. 1459576.

42. Paoletti, P. and J. Neyton, NMDA receptor subunits: function and pharmacology. Current opinion in pharmacology, 2007. 7(1): p. 39–47.

43. Qu, Q., et al., Functional investigation of SLC1A2 variants associated with epilepsy. Cell Death & Disease, 2022. 13(12): p. 1063.

44. Xu, Y., et al., SLC1A2 rs3794087 are associated with susceptibility to Parkinson’s disease, but not essential tremor, amyotrophic lateral sclerosis or multiple system atrophy in a Chinese population. Journal of the neurological sciences, 2016. 365: p. 96–100.

45. Lohr, K.M. and G.W. Miller, VMAT2 and Parkinson’s disease: harnessing the dopamine vesicle. Expert review of neurotherapeutics, 2014. 14(10): p. 1115–1117.

46. Zhai, D., et al., SLC6A3 is a risk factor for Parkinson’s disease: A meta-analysis of sixteen years’ studies. Neuroscience letters, 2014. 564: p. 99–104.

47. Ma, J., et al., Prion-like mechanisms in Parkinson’s disease. Frontiers in neuroscience, 2019. 13: p. 552.

