## Supplementary figures for "Multiomic Spatial Imaging Assay (MSIA) – A High-plex *In Situ* Detection Method for mRNAs, Proteins and Protein-Protein Interactions using Manual And Semi-Automated Workflows"

Advanced Cell Diagnostics (ACD)/Bio-techne, Research & Development, Newark, CA, USA

*Authors contributed equally to the work.

Corresponding Authors:

Chengxin Zhou;

Li-Chong Wang

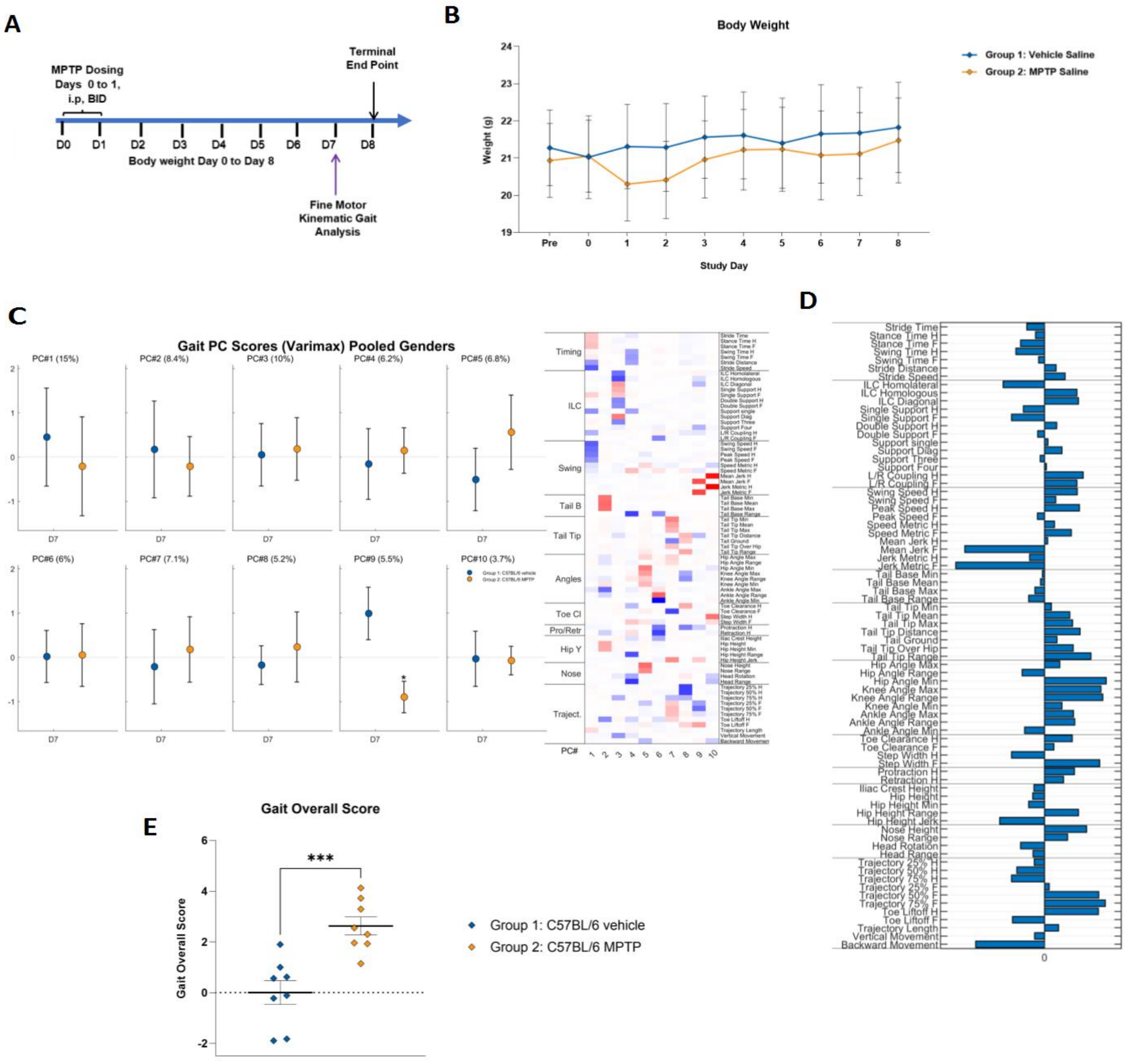


**Supplementary Figure S1**: Establishing MPTP Mouse Model (A) Schematic of study paradigm. (B)Body weight of vehicle or MPTP injected mice. Data are presented as mean ± SEM. Group 1: Sham (n=8; 4 males and 4 females), Group 2: MPTP + Vehicle (n=8; 4 males and 4 females), No statistical significance observed when comparing Group 1 vs. Group 2 (Two-way RM ANOVA). (C) Gait characteristics and fine motor skills of C57BL/6 mice during walking presented as principal component score by PCA analysis. Data are presented as mean ± SEM. Group 1: Vehicle (n=8), Group 2: MPTP (n=8). Left: Corresponding PC Scores (%variance). Higher PC score means: PC#1: decrease of speed; PC#2: overall hip height; PC#3: diagonal gait; PC#4: decrease of vertical hip movement; PC#5: increased knee angles and nose height; PC#6: knee angle and protraction/retraction; PC#7: tail tip height; PC#8: decrease of hindlimb trajectory profile; PC#9: decrease of forelimb trajectory; PC#10: step width and hind limb movement jerkiness. Right: Heat map indicates which mutually correlated parameters are loaded in each ten PCs. Red color means positive correlation, while blue means negative and white no correlation. (D) Identified MPTP model-specific pattern of gait parameters that reflects MPTP induced phenotypes. The vertical midline represents Sham (group 1). Longer bar length indicates bigger differences in MPTP model and was more emphasized in Gait Over Score.

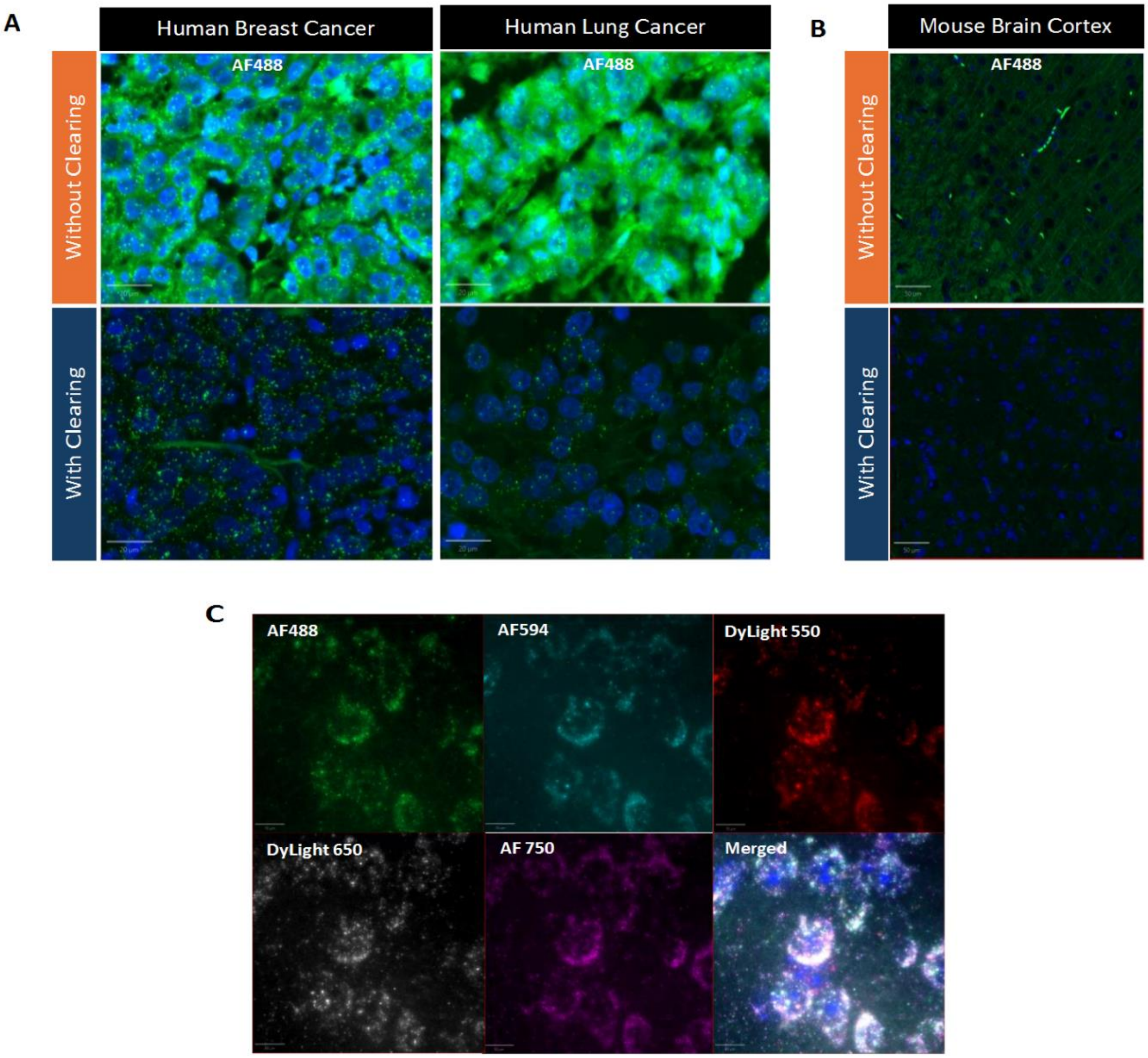


**Supplementary Figure S2**: Result of enzymatic tissue clearing to remove autofluorescence and proteins. Autofluorescence is a major challenge in FFPE tissue analysis, as its intensity can rival that of true fluorescent signals. When autofluorescence is comparable in brightness, it complicates accurate dot detection and quantification, leading to inflated signal counts and false positives (**Supplementary Figure 2A**, top row). This interference compromises decoding accuracy and undermines the reliability of MSIA. To provide a solution to tissues with high autofluorescence, we developed a tissue clearing protocol based on previous publications that can significantly reduce autofluorescence without compromising positive signals. We first tested this method in FFPE human breast cancer and lung cancer tissues using RNAscope HiPlex assay. Without tissue clearing, both tissues have high autofluorescence, and it is very challenging to tell POLR2A signals from autofluorescence. After tissue clearing, both tissues showed significantly lower autofluorescence, and POLR2A signals can be easily picked up and quantified (**Supplementary Figure 2A**, bottom row). Similar results can be observed in FFPE mouse brain (**Supplementary Figure 2B**). We then applied the 100-plex MSIA panel after tissue clearing. All channels showed sharp and punctate signals without interference from autofluorescence (**Supplementary Figure 2C**). These data showed that our method can effectively lower autofluorescence in various FFPE tissues, and it is fully compatible with the MSIA assay.

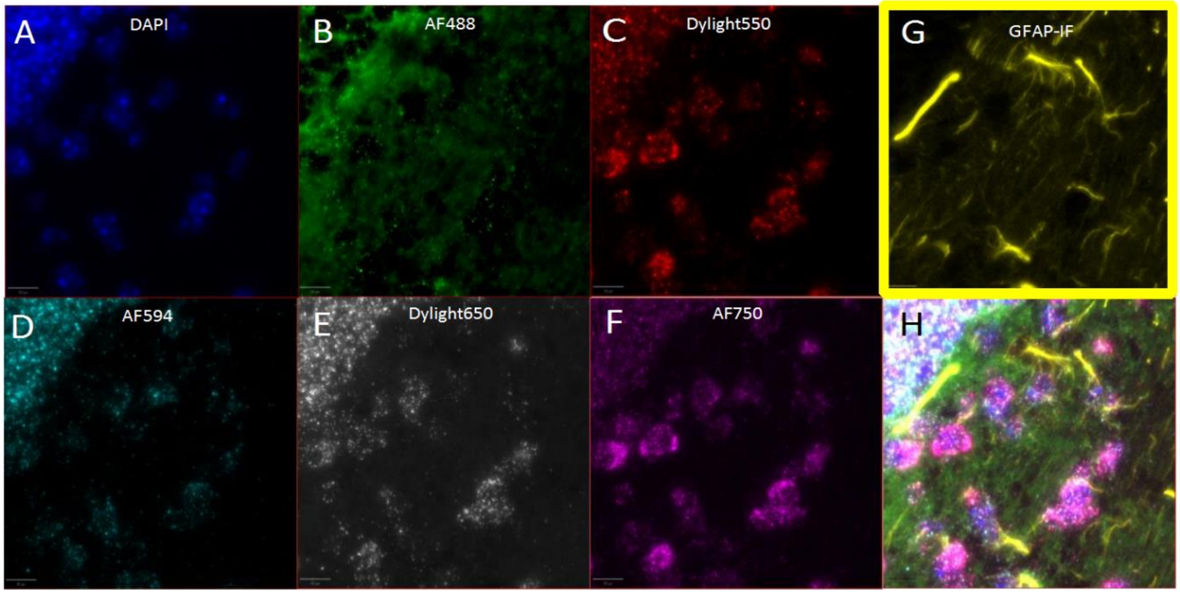


**Supplementary Figure S3**: Overlay of RNA and protein co-detection in MSIA assay.

(A-G) Representative images of 3-cycle MSIA assay with same-slide RNA and GFAP protein co-detections. By using a protease-free tissue pretreatment condition, MSIA enabled 53 RNA transcriptomic profiling on FFPE mouse brain within cycle-1 and cycle-2. Following removal of ISH signals in the tissue slide, GFAP protein immunofluorescence staining was performed in cycle-3 to visualize astrocyte population in the same brain region. Individual ISH channels from 1$^{st}$ cycle of MSIA assay (**A-F**) and GFAP staining from cycle-3 (**G**) in mouse cerebellum are shown. (**H**) Overlay of all 7 signal channels.

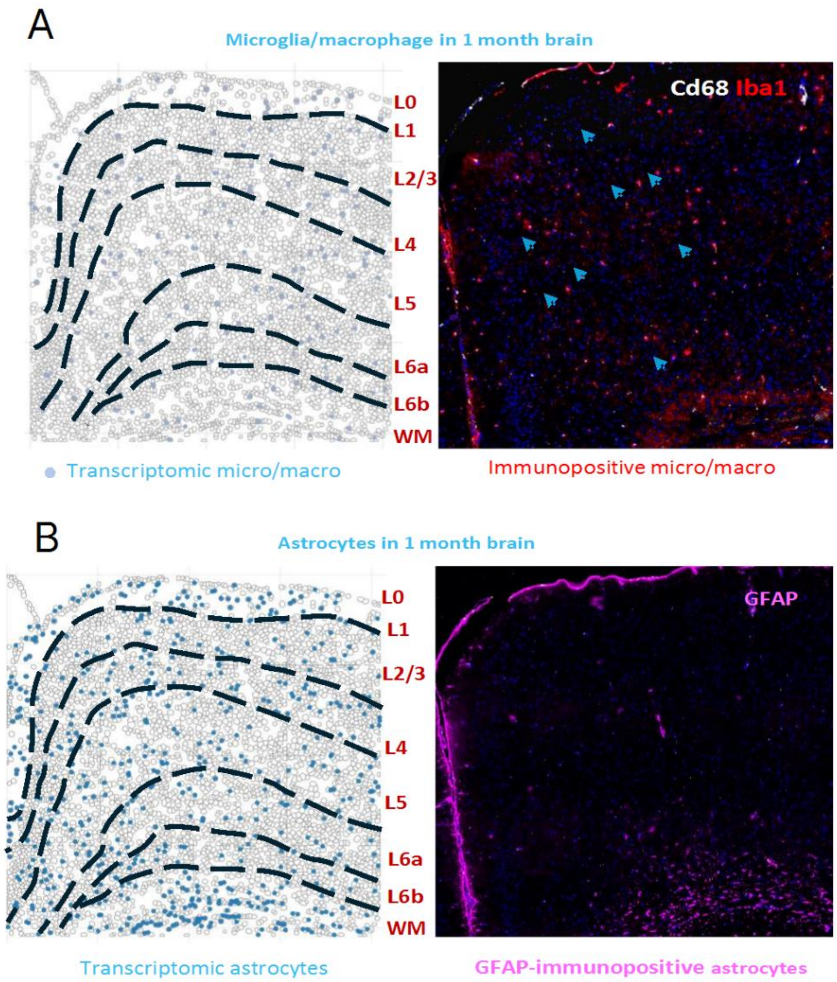


**Supplementary Figure S4**: Mouse brain cell in situ mapping with MSIA spatial transcriptomic data. Single-cell spatial transcriptomics data from ACD-100-plex is mapped into a reduced dimensionality space, using PCA, and the closest match between ACD-100-plex clusters and scRNA-seq taxonomic groups were determined using cosine distance. This approach helps integrate data from these two different modalities by identifying corresponding cell types or clusters between the ACD-100-plex spatial data and the scRNA-seq data. A hierarchical clustering approach is performed to generate the desired set of cell-type-cluster: Firstly, cells are clustered into neurons and non-neuronal cells. Secondly, the neurons are classified into excitatory and inhibitory neurons (MSNs and non-MSNs). Thirdly, the non-neuronal cells are clustered into astrocytes, oligodendrocytes, OPCs, microglia, vascular cells, and immune cells. (**A**) *Left*: spatial representation of MSIA microglial/tissue-resident macrophages in 1-month-old mouse brain cortex (MOp). *Right*: immunofluorescent staining micrograph of the MOp region using primary antibodies against mouse Cd68 and Iba1. (**B**) *Left*: spatial representation of MSIA astrocyte population in 1-month-old mouse brain cortex (Mop).

*Right*: immunofluorescent staining micrograph of the MOp region using primary antibodies against mouse GFAP.

| Genes | cycle-1 barcode | | | | | cycle-2 barcode | | | | |
|---|---|---|---|---|---|---|---|---|---|---|
| | AF488 | Dy550 | Dy594 | Dy650 | AF750 | AF488 | Dy550 | Dy594 | Dy650 | AF750 |
| 2010300C02Rik | 0 | 0 | 0 | 1 | 1 | 1 | 0 | 1 | 0 | 0 |
| Arhgef28 | 0 | 0 | 0 | 1 | 1 | 1 | 1 | 0 | 0 | 0 |
| Bcl11b | 0 | 0 | 1 | 0 | 1 | 1 | 0 | 1 | 0 | 0 |
| Bhlhe22 | 0 | 0 | 1 | 0 | 1 | 1 | 1 | 0 | 0 | 0 |
| Cabp7 | 0 | 0 | 1 | 1 | 0 | 1 | 0 | 1 | 0 | 0 |
| Cpne4 | 0 | 0 | 1 | 1 | 0 | 1 | 1 | 0 | 0 | 0 |
| Necab2 | 0 | 1 | 0 | 0 | 1 | 1 | 0 | 1 | 0 | 0 |
| Prdm8 | 0 | 1 | 0 | 0 | 1 | 1 | 1 | 0 | 0 | 0 |
| Syndig1 | 0 | 1 | 0 | 1 | 0 | 1 | 0 | 1 | 0 | 0 |
| Cpne6 | 0 | 1 | 0 | 1 | 0 | 1 | 1 | 0 | 0 | 0 |
| Epha4 | 0 | 1 | 1 | 0 | 0 | 1 | 0 | 1 | 0 | 0 |
| Hat1 | 0 | 1 | 1 | 0 | 0 | 1 | 1 | 0 | 0 | 0 |
| Neurod6 | 1 | 0 | 0 | 1 | 0 | 0 | 1 | 1 | 0 | 0 |
| Npy2r | 1 | 0 | 0 | 1 | 0 | 1 | 0 | 0 | 0 | 1 |
| Nrp2 | 1 | 0 | 0 | 1 | 0 | 1 | 0 | 0 | 1 | 0 |
| Shisa6 | 1 | 0 | 0 | 1 | 0 | 1 | 0 | 1 | 0 | 0 |
| Pcsk5 | 1 | 0 | 0 | 1 | 0 | 1 | 1 | 0 | 0 | 0 |
| Satb2 | 1 | 0 | 0 | 0 | 1 | 0 | 0 | 0 | 1 | 1 |
| Tle4 | 1 | 0 | 0 | 0 | 1 | 0 | 0 | 1 | 0 | 1 |
| Tox | 1 | 0 | 0 | 0 | 1 | 0 | 0 | 1 | 1 | 0 |
| Cntn6 | 1 | 0 | 0 | 0 | 1 | 0 | 1 | 0 | 0 | 1 |
| Nxph3 | 1 | 0 | 0 | 0 | 1 | 0 | 1 | 0 | 1 | 0 |
| Sema5b | 1 | 0 | 0 | 0 | 1 | 0 | 1 | 1 | 0 | 0 |
| Stard5 | 1 | 0 | 0 | 0 | 1 | 1 | 0 | 0 | 0 | 1 |
| Vwc2l | 1 | 0 | 0 | 0 | 1 | 1 | 0 | 0 | 1 | 0 |
| Col19a1 | 1 | 0 | 0 | 0 | 1 | 1 | 0 | 1 | 0 | 0 |
| Necab1 | 1 | 0 | 0 | 0 | 1 | 1 | 1 | 0 | 0 | 0 |
| Slc44a5 | 1 | 0 | 1 | 0 | 0 | 0 | 0 | 0 | 1 | 1 |
| Calb1 | 1 | 0 | 1 | 0 | 0 | 0 | 0 | 1 | 0 | 1 |
| Lypd6 | 1 | 0 | 1 | 0 | 0 | 0 | 0 | 1 | 1 | 0 |
| Pdyn | 1 | 0 | 1 | 0 | 0 | 0 | 1 | 0 | 0 | 1 |
| Rab3b | 1 | 0 | 1 | 0 | 0 | 0 | 1 | 0 | 1 | 0 |
| Adamtsl1 | 1 | 0 | 1 | 0 | 0 | 0 | 1 | 1 | 0 | 0 |
| Prss35 | 1 | 0 | 1 | 0 | 0 | 1 | 0 | 0 | 0 | 1 |
| Zfpm2 | 1 | 0 | 1 | 0 | 0 | 1 | 0 | 0 | 1 | 0 |
| Fgd5 | 1 | 0 | 1 | 0 | 0 | 1 | 0 | 1 | 0 | 0 |
| Bdnf | 1 | 0 | 1 | 0 | 0 | 1 | 1 | 0 | 0 | 0 |
| Cux2 | 1 | 1 | 0 | 0 | 0 | 0 | 0 | 0 | 1 | 1 |
| Mdga1 | 1 | 1 | 0 | 0 | 0 | 0 | 0 | 1 | 0 | 1 |
| Tmem132d | 1 | 1 | 0 | 0 | 0 | 0 | 0 | 1 | 1 | 0 |
| Dpyd | 1 | 1 | 0 | 0 | 0 | 0 | 1 | 0 | 0 | 1 |
| Pkib | 1 | 1 | 0 | 0 | 0 | 0 | 1 | 0 | 1 | 0 |
| Adamts2 | 1 | 1 | 0 | 0 | 0 | 0 | 1 | 1 | 0 | 0 |
| Cbln4 | 1 | 1 | 0 | 0 | 0 | 1 | 0 | 0 | 0 | 1 |
| Gsg1l | 1 | 1 | 0 | 0 | 0 | 1 | 0 | 0 | 1 | 0 |
| Nwd2 | 1 | 1 | 0 | 0 | 0 | 1 | 0 | 1 | 0 | 0 |
| Rspo2 | 1 | 1 | 0 | 0 | 0 | 1 | 1 | 0 | 0 | 0 |
| Acsbg1 | 0 | 0 | 0 | 1 | 1 | 0 | 0 | 0 | 1 | 1 |
| Cdh20 | 0 | 0 | 0 | 1 | 1 | 0 | 0 | 1 | 0 | 1 |
| Gfap | 0 | 0 | 0 | 1 | 1 | 0 | 0 | 1 | 1 | 0 |
| Gli3 | 0 | 0 | 0 | 1 | 1 | 0 | 1 | 0 | 0 | 1 |
| Ntsr2 | 0 | 0 | 0 | 1 | 1 | 0 | 1 | 0 | 1 | 0 |
| Rorb | 0 | 0 | 0 | 1 | 1 | 0 | 1 | 1 | 0 | 0 |
| Slc39a12 | 0 | 0 | 0 | 1 | 1 | 1 | 0 | 0 | 0 | 1 |
| Cacna2d2 | 0 | 0 | 0 | 1 | 1 | 1 | 0 | 0 | 1 | 0 |
| Calb2 | 0 | 0 | 1 | 0 | 1 | 0 | 0 | 0 | 1 | 1 |
| Cdh4 | 0 | 0 | 1 | 0 | 1 | 0 | 0 | 1 | 0 | 1 |
| Ebf3 | 0 | 0 | 1 | 0 | 1 | 0 | 0 | 1 | 1 | 0 |
| Kctd8 | 0 | 0 | 1 | 0 | 1 | 0 | 1 | 0 | 0 | 1 |
| Slc17a6 | 0 | 0 | 1 | 0 | 1 | 0 | 1 | 0 | 1 | 0 |
| Tmem163 | 0 | 0 | 1 | 0 | 1 | 0 | 1 | 1 | 0 | 0 |
| Dner | 0 | 0 | 1 | 0 | 1 | 1 | 0 | 0 | 0 | 1 |
| Gad2 | 0 | 0 | 1 | 0 | 1 | 1 | 0 | 0 | 1 | 0 |
| Hapln1 | 0 | 0 | 1 | 1 | 0 | 0 | 0 | 0 | 1 | 1 |
| Lamp5 | 0 | 0 | 1 | 1 | 0 | 0 | 0 | 1 | 0 | 1 |
| Rasgrf2 | 0 | 0 | 1 | 1 | 0 | 0 | 0 | 1 | 1 | 0 |
| Cd53 | 0 | 0 | 1 | 1 | 0 | 0 | 1 | 0 | 0 | 1 |
| Cd68 | 0 | 0 | 1 | 1 | 0 | 0 | 1 | 0 | 1 | 0 |
| Laptm5 | 0 | 0 | 1 | 1 | 0 | 0 | 1 | 1 | 0 | 0 |
| Siglech | 0 | 0 | 1 | 1 | 0 | 1 | 0 | 0 | 0 | 1 |
| Sla | 0 | 0 | 1 | 1 | 0 | 1 | 0 | 0 | 1 | 0 |
| Trem2 | 0 | 1 | 0 | 0 | 1 | 0 | 0 | 0 | 1 | 1 |
| Gjc3 | 0 | 1 | 0 | 0 | 1 | 0 | 0 | 1 | 0 | 1 |
| Gpr17 | 0 | 1 | 0 | 0 | 1 | 0 | 0 | 1 | 1 | 0 |
| Opalin | 0 | 1 | 0 | 0 | 1 | 0 | 1 | 0 | 0 | 1 |
| Sox10 | 0 | 1 | 0 | 0 | 1 | 0 | 1 | 0 | 1 | 0 |
| Zfp536 | 0 | 1 | 0 | 0 | 1 | 0 | 1 | 1 | 0 | 0 |
| Acta2 | 0 | 1 | 0 | 0 | 1 | 1 | 0 | 0 | 0 | 1 |
| Ano1 | 0 | 1 | 0 | 0 | 1 | 1 | 0 | 0 | 1 | 0 |
| Carmn | 0 | 1 | 0 | 1 | 0 | 0 | 0 | 0 | 1 | 1 |
| Gucy1a1 | 0 | 1 | 0 | 1 | 0 | 0 | 0 | 1 | 0 | 1 |
| Inpp4b | 0 | 1 | 0 | 1 | 0 | 0 | 0 | 1 | 1 | 0 |
| Pln | 0 | 1 | 0 | 1 | 0 | 0 | 1 | 0 | 0 | 1 |
| Sncg | 0 | 1 | 0 | 1 | 0 | 0 | 1 | 0 | 1 | 0 |
| Aldh1a2 | 0 | 1 | 0 | 1 | 0 | 0 | 1 | 1 | 0 | 0 |
| Col1a1 | 0 | 1 | 0 | 1 | 0 | 1 | 0 | 0 | 0 | 1 |
| Cyp1b1 | 0 | 1 | 0 | 1 | 0 | 1 | 0 | 0 | 1 | 0 |
| Dcn | 0 | 1 | 1 | 0 | 0 | 0 | 0 | 0 | 1 | 1 |
| Fmod | 0 | 1 | 1 | 0 | 0 | 0 | 0 | 1 | 0 | 1 |
| Gjb2 | 0 | 1 | 1 | 0 | 0 | 0 | 0 | 1 | 1 | 0 |
| Igf2 | 0 | 1 | 1 | 0 | 0 | 0 | 1 | 0 | 0 | 1 |
| Spp1 | 0 | 1 | 1 | 0 | 0 | 0 | 1 | 0 | 1 | 0 |
| Grik3 | 0 | 1 | 1 | 0 | 0 | 0 | 1 | 1 | 0 | 0 |
| Htr1f | 0 | 1 | 1 | 0 | 0 | 1 | 0 | 0 | 0 | 1 |
| Meis2 | 0 | 1 | 1 | 0 | 0 | 1 | 0 | 0 | 1 | 0 |
| Myo16 | 1 | 0 | 0 | 1 | 0 | 0 | 0 | 0 | 1 | 1 |
| Sema3e | 1 | 0 | 0 | 1 | 0 | 0 | 0 | 1 | 0 | 1 |
| Slc6a3 | 1 | 0 | 0 | 1 | 0 | 0 | 0 | 1 | 1 | 0 |
| Syt6 | 1 | 0 | 0 | 1 | 0 | 0 | 1 | 0 | 0 | 1 |
| Th | 1 | 0 | 0 | 1 | 0 | 0 | 1 | 0 | 1 | 0 |

**Supplementary Table 1**: a comprehensive list of the mouse gene targets in the 100plex MSIA mouse brain aging panel and the pre-assigned 10-bit barcodes.

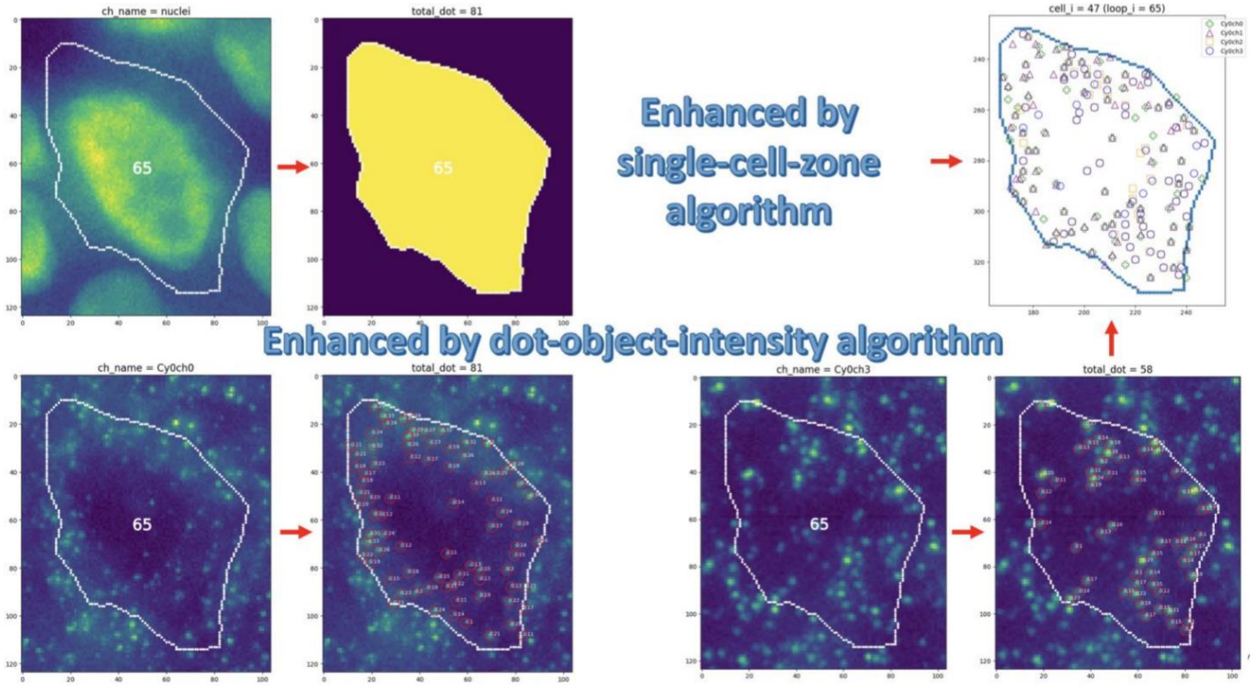


**Supplementary Figure S5: ACDnet signal detections within single-cell-zone.**

Weak RNA signals are detected by a dedicated dot-object-intensity algorithm which focuses on two criteria: the dot prediction from machine-learning model and the intensity value of the dot surrounding area.

Further, only RNA signals within each single cell zone are evaluated: noise outside the cell zone is compressed by a dedicated single-cell-zone algorithm.

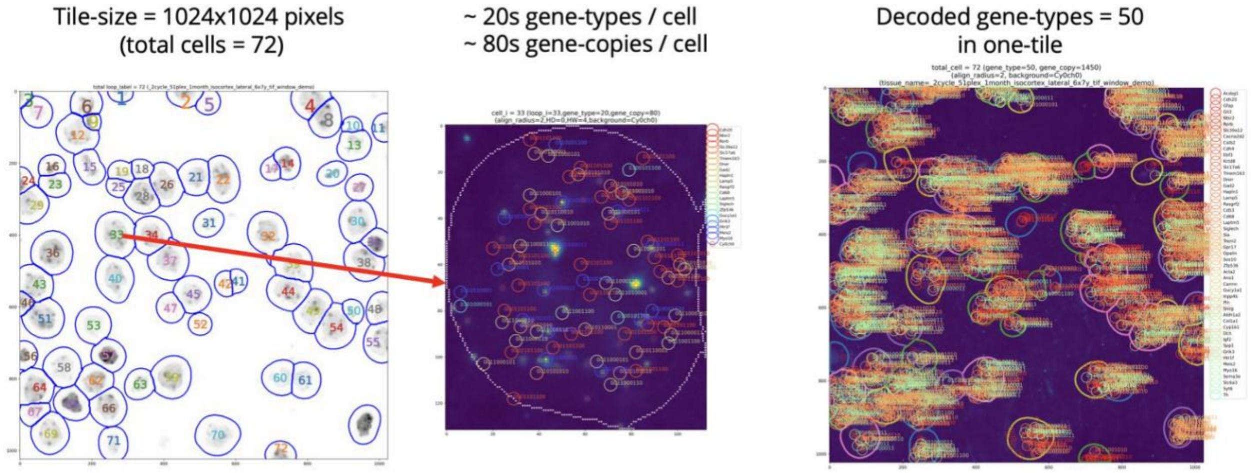


**Supplementary Figure S6: ACDnet cell segmentation characterizations within single-tile.**

(A) left graph: this demo image tile is segmented into 72 single-cell-zones. (B) middle graph: the RNA signals within the demo single-cell-zone are detected into around 80 gene-copies in which 20 gene-types are decoded. (C) right graph, a total of 50 gene-types is decoded in this demo image tile.
